# KIF2A maintains cytokinesis in mouse embryonic stem cells by stabilising intercellular bridge microtubules

**DOI:** 10.1101/2024.07.11.603034

**Authors:** Lieke Stockmann, Hélène Kabbech, Gert-Jan Kremers, Brent van Herk, Bas Dille, Mirjam van den Hout, Wilfred F.J. van IJcken, Dick Dekkers, Jeroen A.A. Demmers, Ihor Smal, Danny Huylebroeck, Sreya Basu, Niels Galjart

## Abstract

Cytokinesis, the final stage of cell division, serves to physically separate daughter cells while ensuring correct segregation of cellular components. In cultured naïve mouse embryonic stem cells cytokinesis lasts unusually long but the underlying mechanisms are not well understood. Here, using cellular and *in vitro* approaches, we describe a novel function for the kinesin-13 member KIF2A in this process. In genome-engineered mouse embryonic stem cells we find that KIF2A mainly localises to spindle poles during metaphase and regulates spindle length in a manner consistent with its known role as microtubule minus-end depolymerase. By contrast, during cytokinesis we observe tight binding of KIF2A on the lattices of intercellular bridge microtubules. At this stage KIF2A maintains microtubule length and number, and controls microtubule acetylation. Based on *in vitro* experiments we propose that the conversion of KIF2A from a depolymerase to a stabiliser is driven both by the inhibition of its ATPase activity, which increases affinity for the lattice, and by a preference of KIF2A for compacted lattices. We propose that during cytokinesis KIF2A maintains the compacted microtubule state, thereby dampening acetylation. As KIF2A depletion causes pluripotency problems and affects mRNA homeostasis our results furthermore indicate that KIF2A-mediated microtubule stabilisation prolongs cytokinesis to maintain pluripotency.

## Introduction

Microtubules (MTs) are dynamic filaments that are part of the cytoskeleton and that are involved in many cellular processes, including cell division, differentiation and migration (Goodson and Jonasson, 2018). MTs are hollow tubes made up of α- and β-tubulin heterodimers, which are arranged in a head-to-tail manner into protofilaments, thirteen of which fold up to form a MT. This arrangement causes an intrinsic MT polarity, exposing β-tubulin at one end (called the plus-end) and α-tubulin at the other (the minus-end). Tubulin dimers assemble into MTs in a GTP-bound form, and GTP hydrolysis occurs once dimers are incorporated in the lattice. Isolated MTs exhibit growth and shrinkage at both ends, as well as conversion from growth to shrinkage (catastrophe) or from shrinkage to growth (rescue). The rate of growth is higher at the plus-end resulting in a “GTP cap” arising from the excess of unhydrolysed GTP. It has been proposed that GTP hydrolysis in MTs is accompanied by changes in the longitudinal distance between dimers in a protofilament (interdimer distance), with GDP-MT lattices displaying a compacted state and MTs made up of the slowly hydrolysable GTP analog GMPCPP, which mimics the GTP state at growing MT ends, having an expanded conformation (Zhang et al., 2018).

The dynamic behaviour of MTs is modulated both by posttranslational modifications (PTMs), as well as by MT associated proteins (MAPs). Acetylation at Lysine 40 (K40) of alpha-tubulin is a long-known PTM that occurs in the lumen of the MT (Janke and Montagnac, 2017). MT acetylation levels increase as MTs age, and acetylation appears to protect MTs from mechanical wear down and breakage (Portran et al., 2017). In order to acetylate MTs, αTAT1 (the enzyme that acetylates alpha-tubulin) has to enter the lumen of ICB MTs to access the lumenal K40. Interestingly, αTAT1 has been shown to act preferentially on MTs with an expanded MT lattice (Shen and Ori-McKenney, 2024).

MAPs are categorised into subclasses based on MT localisation pattern or on structural or functional similarity. Examples of the first category are the plus-end tracking proteins (+TIPs), which accumulate at the plus-ends of growing MTs (Galjart, 2010), and the minus-end targeting proteins (-TIPs), which accumulate at MT minus-ends (Akhmanova and Steinmetz, 2019). An example of the second category is the kinesin superfamily, consisting of MAPs with an ATPase/motor domain which is generally used to propel the proteins over MTs in a directional manner, resulting in transport of cargo associated with the kinesins (Hirokawa et al., 2009). Kinesin-13 family members have their motor domain in the middle of the protein instead of within the N- or C-terminal segments, and are atypical as they do not use ATP hydrolysis for motility, but to depolymerise MTs (Desai et al., 1999). There are four kinesin-13 proteins in mammals, i.e. KIF2A, KIF2B, KIF2C (or MCAK) and KIF24 (Walczak et al., 2013).

During mitosis, chromosomes duplicated in S-phase are separated into two newly formed, diploid nuclei of daughter cells (Mitchison, 1989). The mitotic spindle, which is essential for chromosome segregation, is formed in prophase concurrent with chromosome condensation and nuclear envelope breakdown. The highly dynamic spindle consists of two spindle poles from which various types of MTs emanate. Maintenance of spindle size is critical for chromosome congression, the process of aligning chromosomes at the spindle equator during metaphase (Maiato et al., 2017), and is controlled by MT flux, a continuous poleward movement of MTs (Barisic et al., 2021). It has been shown that KIF2A depolymerises MT minus-ends near spindle poles and plays an important role in regulating poleward flux (Rogers et al., 2004; Sun et al., 2021). After the proper attachment of all chromosomes to the spindle and correct alignment of chromosomes the spindle assembly checkpoint (SAC) is satisfied (Rieder and Maiato, 2004) and anaphase commences, during which chromosomes are pulled apart. At the same time, the network of overlapping interpolar MTs connecting the two spindles rearranges to form central spindle or midzone MTs between the segregated chromosomes (Douglas and Mishima, 2010). Subsequently, during telophase, daughter nuclei are formed.

Cytokinesis, the final stage of cell division, divides the contents of a cell into two daughter cells (Green et al., 2012). Cytokinesis is characterised by constriction of the cell membrane and formation of an intercellular bridge (ICB), which contains the network of midzone MTs (henceforth termed ICB MTs) and the midbody or Flemming body, a dense proteinaceous structure. Once chromosomes and intracellular material have been correctly divided over the daughter cells, abscission occurs, which physically separates the daughter cells. During abscission, adaptor proteins such as CEP55 recruit members of the Endosomal Sorting Complex required for Transport (ESCRT)-III machinery which includes for example, the CHMP family of proteins and the ESCRT-III subunit Ist1, eventually leading to membrane constriction and scission (Carlton et al., 2012; Elia et al., 2011; Paine et al., 2023).

Mouse embryonic stem cells (mESCs) are defined by two properties: pluripotency (the ability to give rise to all somatic lineages and the germline), and self-renewal (proliferation while preserving pluripotency) (Kinoshita and Smith, 2018; Martello and Smith, 2014). Whereas in the mouse embryo pluripotent cells exist for approximately one day, self-renewal is essential for pluripotency maintenance in cultured mESCs. Interestingly, cytokinesis takes several hours in mESCs, which is unusually long and has been coupled to pluripotency maintenance (Chaigne et al., 2020). The mechanisms that control cytokinesis duration remain largely undefined.

Here, we investigate the function of KIF2A in pluripotent mESCs. We find that KIF2A has a dual role. Consistent with studies in differentiated cells (Rogers et al., 2004; Sun et al., 2021), KIF2A functions as a depolymerase during metaphase in mESCs. However, we find that KIF2A converts from a depolymerase to a MT stabilising factor during cytokinesis, where it is required for the maintenance of MTs at the ICB and for cytokinesis duration. We propose that KIF2A-mediated microtubule stabilisation prolongs cytokinesis to maintain pluripotency of naïve mESCs.

## Results

### Localisation of GFP-KIF2A in mESCs

To examine the cellular localisation and function of KIF2A we engineered various *Kif2a* knock-in mESC lines using CRISPR-Cas9-based genome-editing. We inserted four consecutive tags just before the stop codon: eGFP, FKBP12^F36V^ (which is part of the dTAG-13 degradation system for rapid proteasomal depletion of target proteins (Nabet et al., 2018)), a cleavage site for TEV protease, and a Strep-tag II (StrepII) for affinity pull-downs (Fig. S1A). We generated homozygous mESC lines producing endogenous KIF2A-eGFP-FKBP12^F36V^-TEV-StrepII (abbreviated as KIF2A-GFTS). In addition, we generated *Kif2a* knock-out (*Kif2a^KO^*) mESCs, by deleting a large part of the *Kif2a* gene, as well as *Kif2a^GTS^*knock-in mESC lines, containing the same tags as *Kif2a^GFTS^* mESCs, but without FKBP12^F36V^ (Fig. S1A). Western blot analysis demonstrated the presence of KIF2A-eGFP or KIF2A-GFTS fusion proteins of the right size, and absence of KIF2A in homozygous *Kif2a^KO^*mESCs (Fig. 1A).

**Figure 1.**
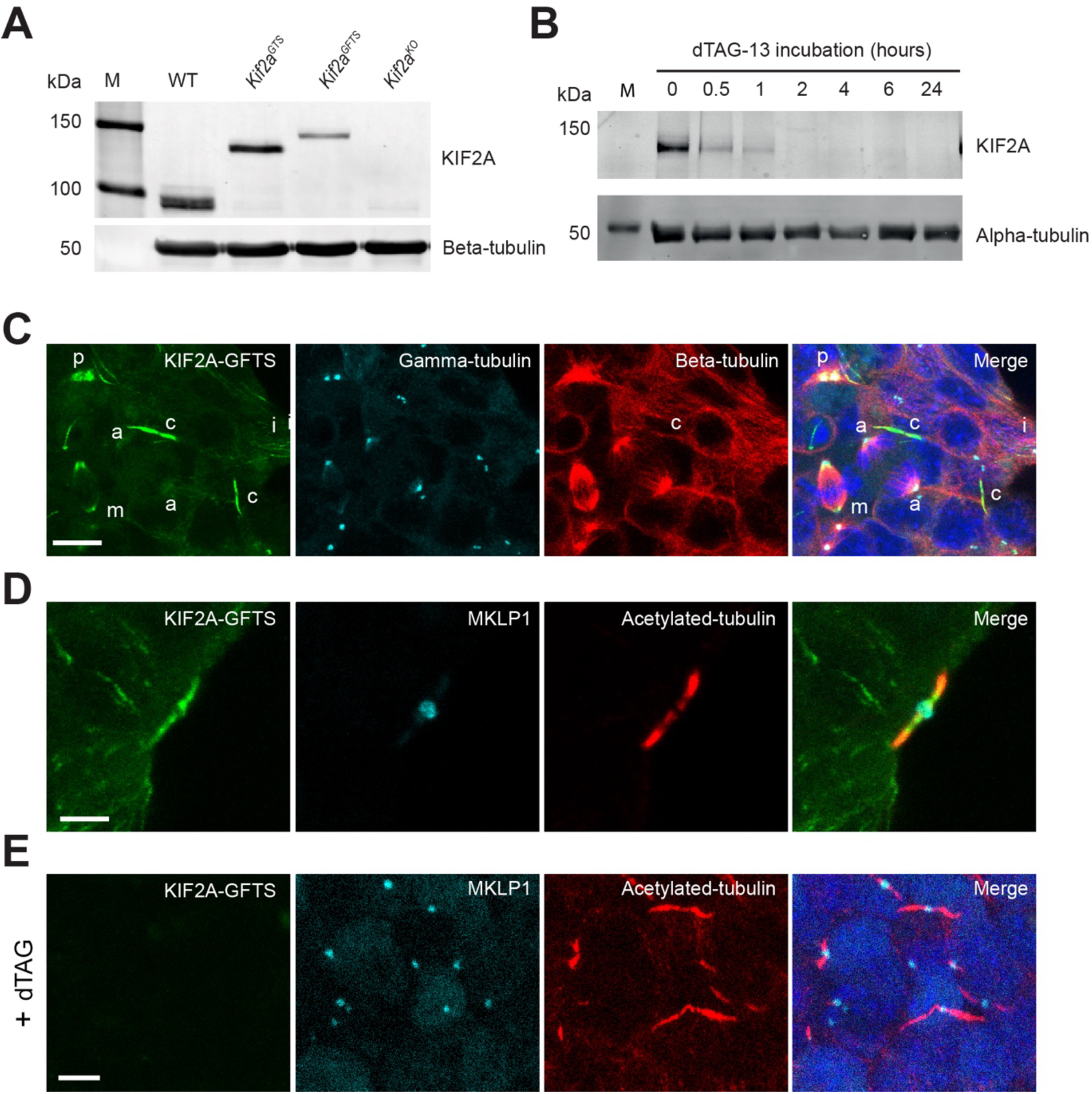
KIF2A accumulates at metaphase spindle poles and ICB MTs. **A**) Genome-edited mESC lines expressing modified or no KIF2A. Lysates of the indicated mESC lines were analysed by western blot using anti-KIF2A (upper panel) or anti-tubulin (lower panel) antibodies. Marker protein lane (M) and relative molecular mass (in kDa) are shown to the left. **B**) Time course-depletion of KIF2A-GFTS. *Kif2a^GFTS^* mESCs were incubated with 50 nM dTAG-13 for the indicated times after which mESC lysates were analysed by western blot with anti-KIF2A (upper panel) or anti-tubulin (lower panel) antibodies. Marker protein lane (M) and relative molecular mass (in kDa) are shown to the left. **C-E**) Intracellular localisation of KIF2A-GFTS. IF analysis of *Kif2a^GFTS^* mESCs stained for KIF2A-GFTS (green) together with anti-gamma-tubulin (cyan) and anti-beta-tubulin (red) antibodies (**C**) or with anti-MKLP1 (cyan), a midbody marker, and anti-acetylated tubulin (red) antibodies (**D, E**). In the merged images DNA (DAPI, blue) is also visualised. In (**E**) dTAG-13 was added to deplete KIF2A and demonstrate specificity of the green signal in IF. In (**C**) mESCs with MT-based structures in the various phases of mitosis are indicated (a: anaphase, m: metaphase, p: prophase), as are ICB MTs (c: cytokinesis). Note that the tubulin signal in ICB MTs is weak because no antigen retrieval was used and the epitope is therefore hidden. Scale bars: 10 μm (**C**), 5 μm (**D, E**).

We noted that in the absence of dTAG-13 the level of KIF2A-GFTS in *Kif2a^GFTS^* mESCs was somewhat lower compared to that of KIF2A in wild-type mESCs (Fig. 1A). Treatment of mESCs with the proteasome inhibitor MG132 did not lead to changes in the level of KIF2A-GFTS (Fig. S1B), indicating that there is no leaky proteasome-mediated degradation of FKBP12^F36V^-containing KIF2A in the absence of dTAG-13. To assess how fast KIF2A-GFTS is degraded in the presence of dTAG-13, we performed a time-course depletion experiment. We could not detect KIF2A on a western blot two hours after addition of dTAG-13 to the medium of mESCs (Fig. 1B). We subsequently treated *Kif2a^GFTS^* mESCs with dTAG-13 for 72 hr. This did not lead to significant cell proliferation defects (Fig. S1C). Thus, in contrast to the essential function of murine KIF2A at birth and during postnatal development (Homma et al., 2003; Homma et al., 2018; Ruiz-Reig et al., 2022), depletion of KIF2A in pluripotent mESCs is not detrimental.

To obtain an overview of KIF2A-GFTS localisation in mESCs, we fixed cells and visualised KIF2A-GFTS together with other structures using specific antibodies (Fig. 1C, D). This immunofluorescence (IF) approach revealed that KIF2A-GFTS was fairly inconspicuous during interphase, although we did observe occasional MT staining. At the onset of mitosis (prophase), KIF2A-GFTS accumulated near spindle pole centrosomes and this localisation was maintained in metaphase (Fig. 1C). In anaphase and telophase KIF2A-GFTS was still near centrosomes, but the signal was weaker (Fig. 1C). During cytokinesis KIF2A-GFTS became highly enriched at the ICB, as shown by co-staining with antibodies against acetylated-tubulin and the midbody marker MKLP1 (Zhu et al., 2005) (Fig. 1C, D). dTAG-13-mediated depletion of KIF2A-GFTS resulted in a complete disappearance of green fluorescence (Fig 1E), demonstrating that the signal observed in non-treated cells represents KIF2A-GFTS. Thus, in mESCs KIF2A-GFTS first accumulates near the spindle poles close to MT minus-ends while later a strong accumulation is observed on ICB MTs, where KIF2A appears to be bound along the length of the MTs, and hence on the lattice of these MTs.

### Dynamic behaviour of KIF2A-GFTS and MTs in pluripotent mESCs

To concurrently analyse the dynamic behaviour of KIF2A-GFTS and MTs we added SiR-Tubulin (Lukinavicius et al., 2014) to mESCs and used light sheet fluorescence microscopy (LSFM) for visualisation, as this method is well suited for long term fluorescence time-lapse imaging of relatively thick 3D samples, and also reduces phototoxicity and photobleaching as compared to confocal microscopy (Tomer et al., 2013). We acquired z-stacks of between 30-60 micron (z-step size of 1 micron), and imaged colonies every 10 minutes for 16 hours.

We detected ICB MTs throughout mESC colonies (examples of a 3D slice-through and 3D projection of KIF2A-GFTS and SiR-tubulin localisation in a mESC colony are shown in Movies S1 (z-stack) and S2 (3D-projection), respectively). In time lapse experiments we observed dynamic behaviour of KIF2A-GFTS and ICB MTs (Fig. 2A, Movie S3). At the onset of cytokinesis, approximately 20 minutes after metaphase, we detected intensely fluorescent midzone MTs (Fig. 2A). KIF2A-GFTS became visible at these MTs approximately 10 minutes later (Fig. 2A). By contrast, at the end of cytokinesis KIF2A-GFTS remained visible longer than SiR-Tubulin (Fig. 2A). Quantification of cytokinesis duration using both fluorescent dyes revealed a longer duration with KIF2A-GFTS compared to SiR-Tubulin (Fig. 2B, duration with KIF2A-GFTS: 6.1 ± 2.2 hr (367 ± 133 min), duration with SiR-Tubulin: 4.9 ± 1.9 hr (292 ± 195 min)). This is shorter than the time (∼8.2 hours) documented previously (Chaigne et al., 2020).

**Figure 2.**
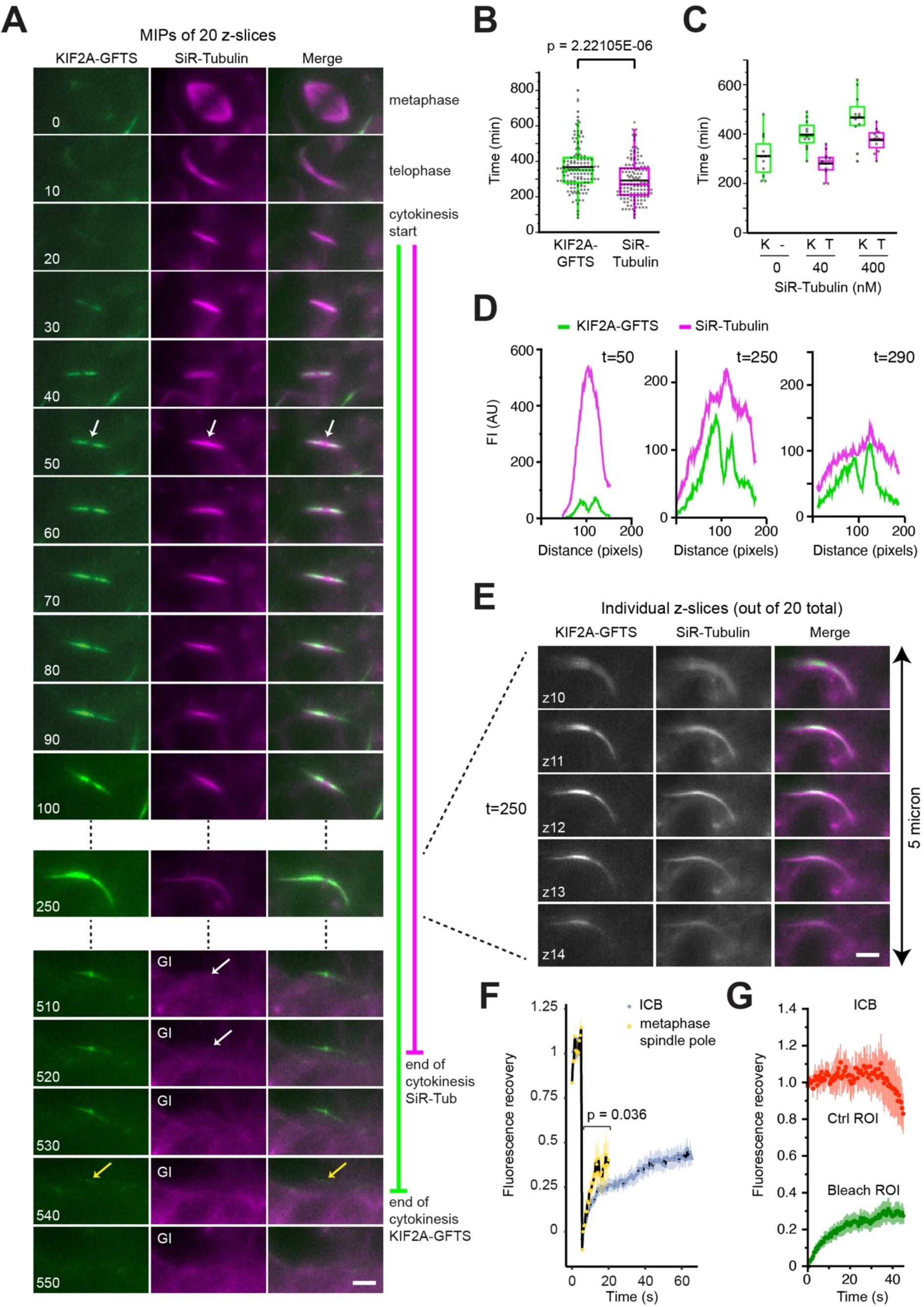
Dynamic localisation of KIF2A-GFTS and MTs during cytokinesis. **A**) Localisation of KIF2A-GFTS and MTs in live mESCs using light sheet fluorescence microscopy (LSFM). We acquired images of KIF2A-GFTS (green) and SiR-tubulin (magenta) every 10 min and designated metaphase as the start point (t=0). We examined 20 z-slices (1 micron apart) at each time point and then generated maximum intensity projections (MIPs) of the individual time points. Selected MIPs are shown with their times indicated in the panels. Start and end of cytokinesis are indicated to the right (end times differ based on fluorescent signal). White arrows at t=50 min indicate MTs that are present in the midbody (the KIF2A-GFTS-negative region) and extend into the “arms” of the ICB. Note that from t=510 onward the SiR-tubulin signal was artificially enhanced, to allow visualisation of remaining ICB MTs (white arrows at t=510 and t=520 min). At t=540 min KIF2A-GFTS is still visible (yellow arrows) albeit barely. Supplementary Video 6 shows the complete time lapse experiment. Scale bar: 4 μm. **B**) Measurement of cytokinesis duration. Whisker plot showing the duration of cytokinesis (in min) in non-treated *Kif2a^GFTS^* mESCs, measured using KIF2A-GFTS (green) or SiR-Tubulin (magenta). Whisker plot shows interquartile range, median (coloured lines), and average (black lines). N = 6 independent experiments, n= 130 ICBs (from 20 mESC colonies). T-test shows significant difference in duration times measured with KIF2A-GFTS or SiR-Tubulin. **C**) Measurement of cytokinesis duration at increasing SiR-Tubulin concentrations. Whisker plot depicts duration of cytokinesis (in min) in wild type mESCs incubated in the indicated SiR-Tubulin concentrations. Duration times were measured using KIF2A-GFTS (K) or SiR-Tubulin (T) (at 0 nM no SiR-Tubulin measurement was possible). Median is indicated with a coloured line, mean with a black line. N=2 independent experiments, n= 12 ICBs (0 nM SiR-Tubulin), 11 ICBs (40 nM SiR-Tubulin), and 11 ICBs (400 nM SiR-Tubulin). **D**) Relocalisation of KIF2A-GFTS and MTs during cytokinesis. The fluorescence intensity (FI) in arbitrary units (AU) of KIF2A-GFTS (green) and SiR-Tubulin (magenta) was plotted along the horizontal axis of ICB MTs (distance in pixels) at t=50, t=250, and t=290. To properly display signals different y-axis scales were used for each plot. **E**) Localisation of KIF2A-GFTS and MTs in individual z-slices of mESC colony. At t=250 KIF2A-GFTS and SiR-Tubulin were detected in 5 consecutive slices (out of the 20). Thus, the ICB MTs span a vertical (z) dimension of 5 micron. Scale bar: 4 μm. **F, G**) Dynamic behaviour of KIF2A-GFTS. In (**F**) a fluorescence recovery after photobleaching (FRAP) experiment is shown, where KIF2A-GFTS was bleached at the metaphase spindle pole (yellow, note that the recovery of KIF2A-GFTS could not be determined beyond 20 seconds, due to the dynamic behaviour of spindle poles, causing movement away from the bleach region), or at ICB MTs (blue). Points depict average of normalised signal and shaded areas depict standard error of mean (SEM). N = 3 independent experiments, n = 10-23 FRAP experiments per location. P-value of the significant difference in slope of the recovery is indicated, as examined with T-test. In (**G**) a FRAP analysis is shown of selected ICB MTs with similar lengths and KIF2A-GFTS distribution. Fluorescence recovery of KIF2A-GFTS on the bleached (Bleach ROI, green) and non-bleached (control (Ctrl) ROI, red) arms of ICB MTs is shown. Points depict average of normalised signal and shaded lines standard error of the mean (SEM). N = 3 independent experiments, n = 13 ICB MTs.

We observed a considerable fluorescence decay of the SiR-tubulin signal during imaging (Fig. S2A), presumably because SiR-Tubulin, which is a taxol-derivative (Lukinavicius et al., 2014), is actively pumped out of mESCs over the course of the imaging experiment. KIF2A-GFTS signal also decreased during the 16 hour time lapse, but to a much lesser extent (Fig. S2B). To enhance the MT signal in live-imaging experiments we increased SiR-tubulin concentration. However, this led to a longer cytokinesis (Fig. 2C). Since SiR-Tubulin is taxol-based and therefore a MT stabiliser, these results suggest that increasing MT stability lengthens cytokinesis duration.

The SiR-Tubulin signal was always detected throughout the ICB, including the midbody, whereas KIF2A-GFTS eluded this central domain until the end of cytokinesis (Fig. 2A, D). We furthermore observed asymmetric accumulation of KIF2A-GFTS and MTs, with one of the future daughter mESCs having more signal than the other (Fig. 2A, D, E, Movie S3). Investigation of fluorescence signal in individual z-sections showed that the asymmetry was not an artefact of maximum intensity projections (Fig. 2E). We examined fluorescence intensities of SiR-Tubulin and KIF2A-GFTS during the early phases of cytokinesis. We detected a 3-fold increase in signal intensity of SiR-Tubulin on spindle and ICB MTs over the cytoplasm, whereas KIF2A-GFTS fluorescence intensity on ICB MTs was approximately 7-fold higher than cytoplasmic fluorescence intensity (Fig. S2C). These data indicate that most KIF2A-GFTS molecules are associated with ICB MTs and that the affinity of KIF2A for ICB MTs is high.

To analyse the dynamic behaviour of KIF2A-GFTS in mESCs we carried out Fluorescence Recovery After Photobleaching (FRAP). We bleached KIF2A-GFTS both at metaphase spindles and on ICB MTs, and observed a faster initial recovery at metaphase spindles compared to ICB MTs (Fig. 2F). At the end of the FRAP experiment, the fraction of fluorescent KIF2A-GFTS molecules on ICB MTs was ∼40% (Fig. 2F). Given the low amount of KIF2A-GFTS present in the cytoplasm (Fig. S2C), exchange of cytoplasmic with MT-bound KIF2A-GFTS is limited. To examine the mechanism underlying the partial recovery of KIF2A-GFTS on ICB MTs, we selected ICB MTs of similar length and KIF2A-GFTS intensity profile, and then analysed fluorescence dynamics on the complete MT structure throughout the FRAP experiment. We observed no obvious fluorescence loss in the non-bleached “arm” (ICB MTs in one of the future daughter mESCs) in the first phase of the recovery (Fig. 2G), indicating there is no diffusion of KIF2A-GFTS from one daughter mESC to the other. Instead, the limited recovery of KIF2A-GFTS that we observed on the bleached arm (Fig. 2F, G) appeared to be partly due to diffusion from the cytoplasmic side of the ICB MTs (Fig. S2D). This diffusion coincided with a decrease of KIF2A fluorescence next to the bleached region (Fig. S2D). These results indicate that KIF2A is tightly bound to the lattice of ICB MTs but can undergo lateral movement within the bundle of MTs. Despite its relatively immobile behaviour in the time window of the FRAP studies (∼1 min), KIF2A-GFTS displays dynamic behaviour within the time frame of the LSFM-based experiments (∼10 min).

### A dual function for KIF2A in mESCs

We next examined the role of KIF2A using KIF2A-depleted mESCs, first focussing on metaphase, and subsequently on cytokinesis. We used two models, i.e. wild-type (WT) versus *Kif2a^KO^* (KO) mESCs, or *Kif2a^GFTS^* mESCs that were either treated for 24 hours with dTAG-13 or not treated. Analysis of fixed mESCs revealed that KIF2A depletion leads to an increased pole-to-pole distance during metaphase in both KIF2A-depletion models (Fig. S3A). Thus, absence of KIF2A increases spindle length, which is consistent with RNAi-based KIF2A knockdown results in other cell types (Gaetz and Kapoor, 2004; Jang et al., 2008; Rogers et al., 2004), and with the notion, supported by *in vitro* experiments (Henkin et al., 2023), that during metaphase KIF2A localises near MT minus-ends on spindle poles and acts as a MT depolymerase.

To investigate cytokinesis, we analysed SiR-tubulin behaviour in control and KIF2A-depleted mESCs. We first ascertained that addition of dTAG-13 to WT mESCs did not affect cytokinesis duration (Fig. 3A). We then found that both *Kif2a^KO^* mESCs (Fig. 3A) and dTAG13-treated *Kif2a^GFTS^*mESCs (Fig. 3B) displayed a significantly shorter cytokinesis duration, compared to wild-type or non-treated *Kif2a^GFTS^* mESCs, respectively. Cytokinesis lasted 5.4 ± 1.1 hr (324 ± 68 min) in wild-type mESCs and 3.8 ± 0.79 hr (227 ± 48 min) in *Kif2a^KO^* mESCs, whereas it lasted 4.9 ± 1.9 hr (292 ± 115 min) in non-treated *Kif2a^GFTS^*mESCs (same SiR-Tubulin data as reported in Fig. 2B), and 4.0 ± 2.1 hr (238 ± 124 min) in dTAG-13-treated *Kif2a^GFTS^* mESCs. Thus, control mESCs display similar cytokinesis duration times, and in the two KIF2A-depletion models the cytokinesis duration is shorter, strongly suggesting that KIF2A regulates cytokinesis in pluripotent mESCs. Notably, the size of the mESC colony did not influence duration times (Fig. 3C).

**Figure 3.**
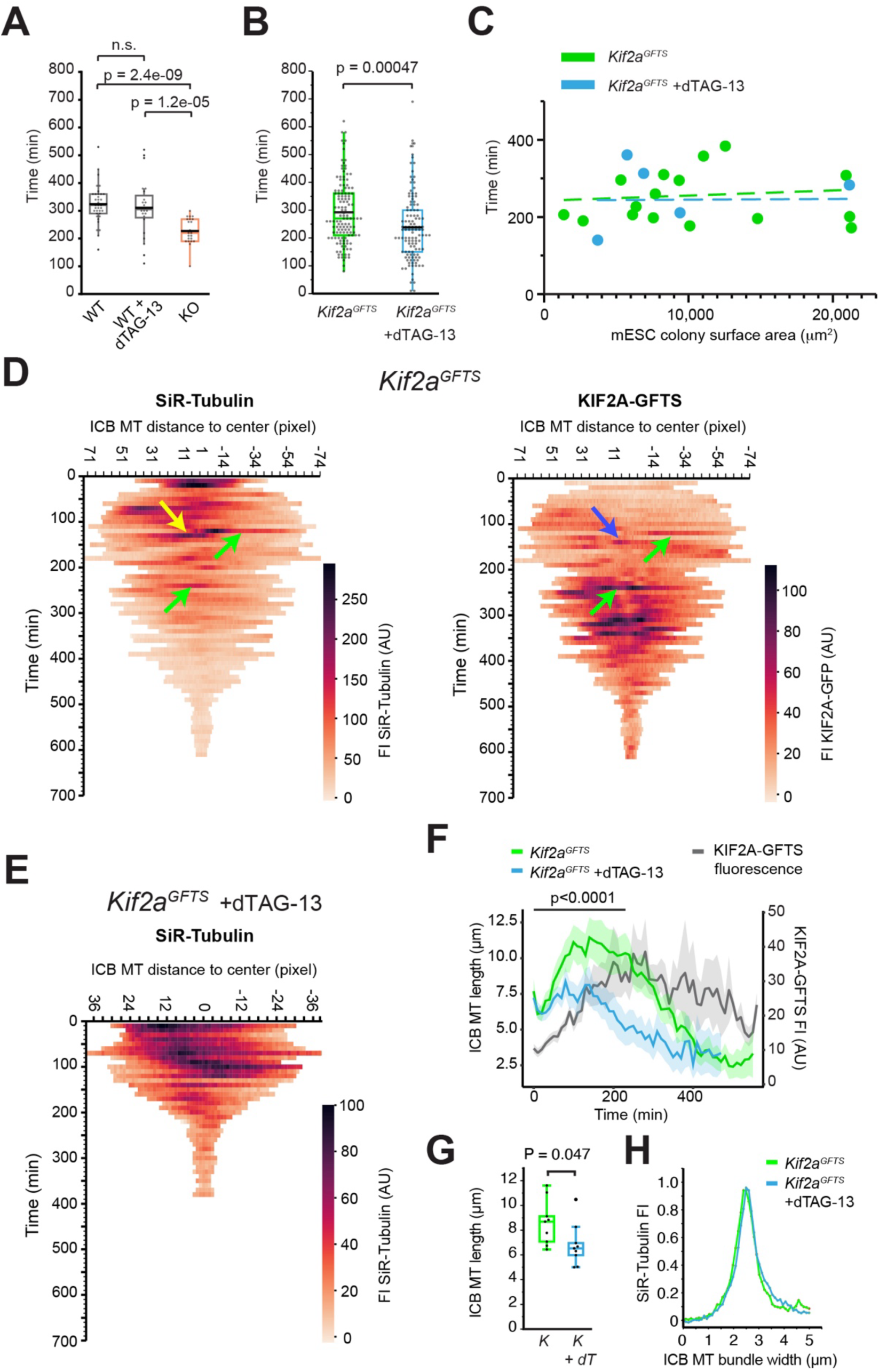
KIF2A regulates cytokinesis duration and ICB MTs. **A, B**) Measurement of cytokinesis duration in control and KIF2A-depleted mESCs. Whisker plot shows the duration of cytokinesis (in min) using SiR-Tubulin signal. Median is indicated with a coloured line, mean with a black line. We measured duration in wild type (WT), WT treated with dTAG-13 and *Kif2a^KO^*(KO) mESCs (**A**) or in non-treated and -dTAG-13-treated *Kif2a^GFTS^*mESCs (**B**). In (**A**) n = 37 (WT), 24 (WT + dTAG-13), 22 (KO), 2 independent experiments. P-values shown from one way ANOVA followed by Tukey’s post-hoc test. In (**B**) n = 130 (*Kif2a^GFTS^*), 118 (*Kif2a^GFTS^* +dTAG-13), 6 and 5 independent experiments, respectively. P-values shown are from T-test. **C)** Measurement of cytokinesis duration in differently sized mESC colonies. Dot plot showing the duration of cytokinesis (in min), measured using SiR-Tubulin in non-treated and -dTAG-13-treated *Kif2a^GFTS^* mESCs. mESC colonies with different surface areas were measured. Stippled lines indicate linear fits. Data are from 6 (*Kif2a^GFTS^*) or 5 (*Kif2a^GFTS^* +dTAG-13) independent experiments. **D, E**) Dynamic behaviour of KIF2A-GFTS and MTs during cytokinesis. Kymographs showing fluorescence intensity (FI, represented as heatmaps) of SiR-tubulin (**D, E**) or KIF2A-GFTS (**D**), over time in non-treated (**D**) or -dTAG-13-treated (**E**) *Kif2a^GFTS^*mESCs. Each horizontal line represents ICB MTs captured at a specific time point. Hence, the length of the line reflects the length of the ICB MTs at that moment in time. Time is represented in 10-minute intervals; heatmap color depicts the local FI of SiR-tubulin or KIF2A-GFTS within the ICB MTs. Green arrows indicate concurrent bouts of high SiR-tubulin and KIF2A-GFTS intensity. Yellow arrow indicates bout of SiR-tubulin only. It is followed 10 min later by an increase in KIF2A-GFTS signal (blue arrow). Notice reduced ICB MT length in dTAG-13-treated *Kif2a^GFTS^* mESC (**E**) compared to non-treated mESCs (**D**, note that different length and heatmap scales were used in (**D**) and (**E**)). Furthermore, cytokinesis lasts considerably shorter in dTAG-13-treated *Kif2a^GFTS^* mESC (**E**). **F**) ICB MT length and KIF2A-GFTS fluorescence over time during cytokinesis. Line plots depicting the ICB MT length in non-treated (green) or dTAG-13-treated (blue) *Kif2a^GFTS^*mESCs (left axis), and of KIF2A-GFTS fluorescence intensity (FI, grey, right axis) in arbitrary units (A.U.), over time. Shaded areas represent the SEM. n = 10 ICBs per condition. P-value derived from repeated measures ANOVA of the difference in ICB length over time between *Kif2a^GFTS^* and dTAG-13-treated *Kif2a^GFTS^* mESCs until 230 minutes. **G**) Length of ICB MTs. The length of ICB MTs was measured 100 minutes after the onset of metaphase, a time point where the effect of KIF2A depletion is maximal (see panel F). Length was measured by tracing SiR-Tubulin fluorescence intensity in non-treated (K) or dTAG-13-treated (K+dT) *Kif2a^GFTS^* mESCs. P-value derived from T-test. **H**) ICB MT bundle thickness at the midbody. SiR-Tubulin fluorescence intensity (FI) of ICB MT bundles at the midbody was measured 100 minutes after the onset of metaphase, a time point where the effect of KIF2A depletion is maximal (see panel F). The thickness was measured by placing a line perpendicular to the SiR-Tubulin-positive MT bundle and measuring FI across the bundle. We measured FI in non-treated (green) or dTAG-13-treated (blue) *Kif2a^GFTS^* mESCs. Data were normalised afterwards.

To analyse the dynamic behaviour of ICB MTs and KIF2A-GFTS in more detail we developed a visualisation tool, which yielded time-length plots (Fig. 3D, E). In these kymographs ICB MTs in each frame of the time lapse movie are represented by a line, and these are plotted in time with an imaging interval of 10 minutes. Fluorescence intensities are represented as heatmaps for each timepoint (Fig. 3D, E, see Movies S4 and S5 for the corresponding time-lapse experiments). Consistent with the data presented in Fig. 2, the kymographs revealed local accumulation of MTs and KIF2A-GFTS within ICB arms. Notably, signals often co-localised within an arm (see green arrows in Fig. 3D), although we also detected independent accumulations (see yellow and blue arrows in Fig. 3D, note that in this example the accumulation of SiR-Tubulin (yellow arrow) is followed 10 min later by an increase in KIF2A-GFTS signal (blue arrow), suggesting positional dependence). By averaging fluorescence intensities across individual kymographs we found that in non-treated *Kif2a^GFTS^* mESCs the ICB MT length increased by about 60% in the first hours of cytokinesis, followed by a gradual decrease afterwards (Fig. 3F).

Thus, cytokinesis in pluripotent mESC has two distinct phases, one of a net ICB MT length increase and one where the ICB MT length decreases. The ICB MT length increase was accompanied by an increase in KIF2A-GFTS signal in time, however, KIF2A-GFTS accumulation continued after the maximum ICB MT peak length was attained, and peaked later (Fig. 3D, F). Strikingly, whereas at the start of cytokinesis the ICB MT length was similar in mESCs lacking KIF2A-GFTS compared to control mESCs, we did not observe a significant ICB MT length increase in KIF2A-depleted mESCs in the first phase of cytokinesis (Fig. 3E, F). We next measured both the length of MT bundles in the ICB, as well as the diameter of these bundles within the midbody region. We examined these parameters in control and KIF2A-depleted mESCs and chose a fixed time point, i.e. 100 minutes after metaphase onset, when the effect of KIF2A depletion is the strongest. Consistent with the kymograph-based results (Fig. 3F) ICB MT length was significantly decreased in KIF2A-depleted mESCs compared to controls 100 minutes after metaphase onset (Fig. 3G). By contrast, the width of ICB MT bundles was not different from control mESCs (Fig. 3H). These results indicate that a similar number of MTs is present in the midbody, where MT plus ends overlap, but that MTs in the arms of KIF2A-depleted mESCs are shorter. In agreement with these data, we observed a similar SiR-Tubulin fluorescence intensity in the midbody region in dTAG-13-treated and non-treated mESCs at the onset of the time lapse experiment (Fig. S3B), yet less SiR-Tubulin throughout the time lapse experiment in ICB MTs of dTAG-13-treated mESCs (Fig. S3C). Collectively, our LSFM-based studies in live mESCs indicate that KIF2A localises in a dynamic fashion along ICB MTs and serves to increase MT length, and not to decrease it, contrary to its expected function as a depolymerase.

### *In vitro* properties of GFP-KIF2A

Early *in vitro* experiments performed with a monomeric “minimal” KIF2A protein containing only the motor domain and the neighbouring neck region, showed that this minimal domain binds to two consecutive tubulins in a protofilament, with the neck of KIF2A binding to one tubulin dimer and the motor domain to the other (Trofimova et al., 2018). This minimal domain was shown to be sufficient to depolymerise MTs (Ogawa et al., 2017; Trofimova et al., 2018). These experiments also revealed that dissociation of KIF2A from MTs is ATP-hydrolysis dependent and accompanied by MT depolymerisation. More recent *in vitro* MT reconstitution assays using purified full length KIF2A revealed that KIF2A is an autonomous -TIP and depolymerase (Henkin et al., 2023).

In an effort to explain our cellular data, in which KIF2A alters its binding depending on mitotic stage, we examined the *in vitro* localisation of KIF2A. We produced and purified human KIF2A fused to eGFP (Fig. S4A), confirmed the purity of the protein by mass spectrometric analysis (Fig. S4B, C, Table S1), and used GFP-KIF2A in *in vitro* MT reconstitution assays. To examine the effect of ATP hydrolysis on GFP-KIF2A binding, we performed experiments both in the presence of ATP, when GFP-KIF2A can dissociate from MTs, as well as in the absence of ATP. At 3 nM GFP-KIF2A + ATP we observed intermittent accumulation of GFP-KIF2A at MT minus-ends (which were distinguished because they were growing slower than the plus-end) and weak accumulation on the MT lattice (Fig. 4A,B, Fig. S5A,B, Movie S6). At 6 nM + ATP we observed a continuous accumulation of GFP-KIF2A at MT minus-ends (Fig. 4C,D). Thus, similar to a recent publication (Henkin et al., 2023), we find that in the presence of ATP GFP-KIF2A is a -TIP, clearly preferring the minus-end over the lattice.

**Figure 4.**
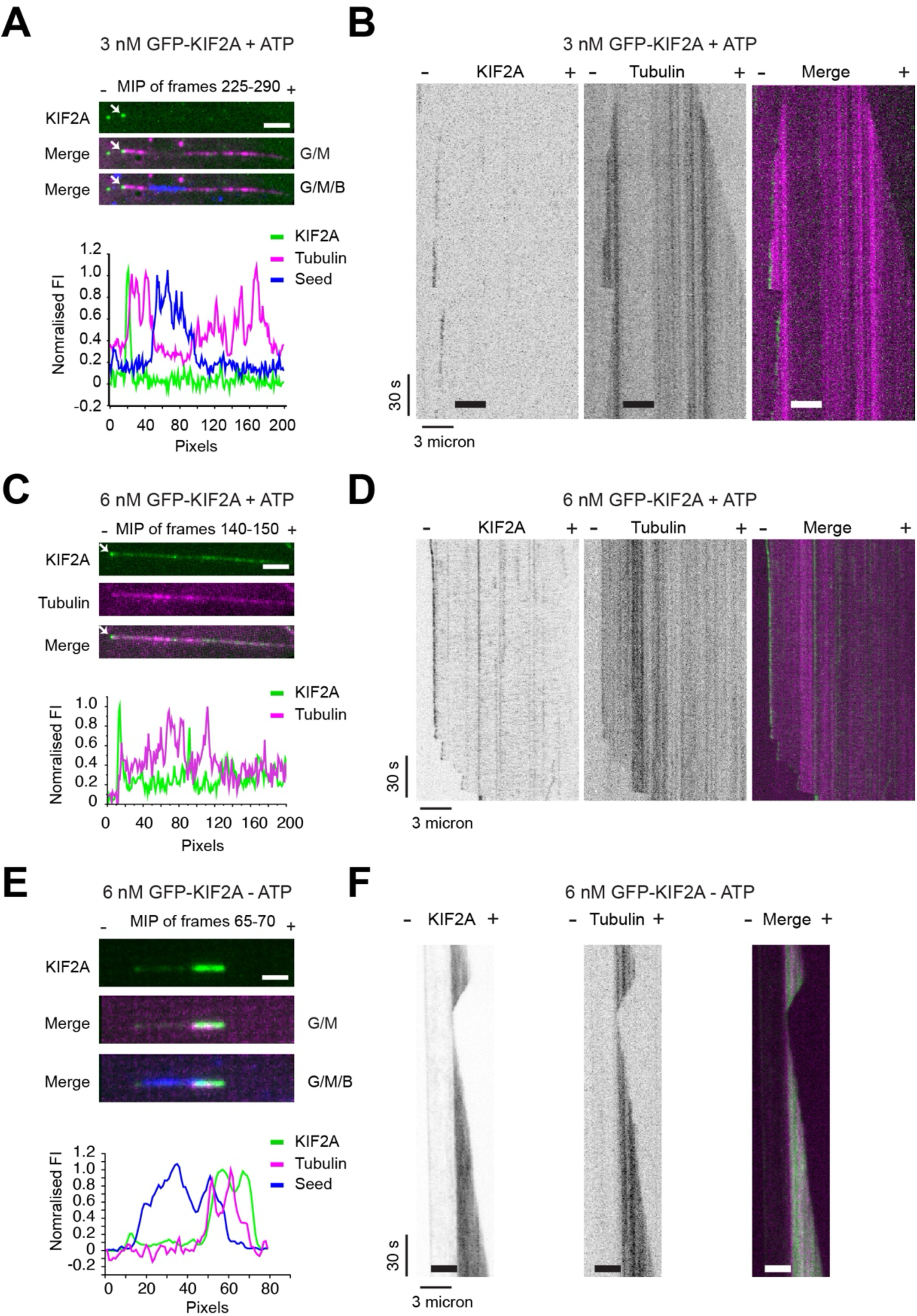
In vitro GFP-KIF2A behaviour. **A**) Maximum intensity projection (MIP) of TIRF microscopy images (frames 225-290) of 3 nM GFP-KIF2A binding to MTs. The experiment was carried out in the presence of ATP. GFP-KIF2A is labeled in green (G) and tubulin in magenta (M). The MT seed is labelled in blue (B). The - and + signs indicate MT minus- and plus-end, respectively. Note that the seed signal was captured in the first frame of the time lapse, and the still image was then superimposed. GFP-KIF2A accumulation on the MT minus-end is indicated by the white arrow. Normalised fluorescence intensity distributions (FI) of the three dyes along the MT are plotted below the MIP. Scale bar: 2 μm. **B**) Kymograph of *in vitro* MT reconstitution assay with 3 nM GFP-KIF2A. The experiment was carried out in the presence of ATP. In the merge, GFP-KIF2A is labeled in green and tubulin in magenta. The MT seed is indicated in the kymographs by a thick line. The - and + signs indicate MT minus- and plus-end, respectively. **C**) Maximum intensity projection (MIP) of TIRF microscopy images (frames 1-2) of 6 nM GFP-KIF2A binding to the MTs. The experiment was carried out in the presence of ATP. GFP-KIF2A is labeled green, tubulin is labeled magenta. GFP-KIF2A accumulation on the MT minus-end is indicated by the white arrow. The - and + signs indicate MT minus- and plus-end, respectively. Normalised fluorescence intensity distributions (FI) of the two dyes along the MT are plotted below the MIP. Scale bar: 2 μm. **D**) Kymograph of *in vitro* MT reconstitution assay with 6 nM GFP-KIF2A. The experiment was carried out in the presence of ATP. In the merge, GFP-KIF2A is labeled in green and tubulin in magenta. The - and + signs indicate MT minus- and plus-end, respectively. **E**) Maximum intensity projection (MIP) of TIRF microscopy images (frames 65-70) of 6 nM GFP-KIF2A binding to MTs. The experiment was carried out in the absence of ATP. GFP-KIF2A is labeled in green (G) and tubulin in magenta (M). The MT seed is labelled in blue (B). The - and + signs indicate MT minus- and plus-end, respectively. Note that the seed signal was captured in the first frame of the time lapse, and the still image was then superimposed. Normalised fluorescence intensity distributions (FI) of the three dyes along the MT are plotted below the MIP. Scale bar: 1 μm. **F**) Kymograph of *in vitro* MT reconstitution assay with 6 nM GFP-KIF2A. The experiment was carried out in the absence of ATP. In the merge, GFP-KIF2A is labeled in green and tubulin in magenta. The MT seed is indicated in the kymographs by a thick line. The - and + signs indicate MT minus- and plus-end, respectively.

Interestingly, when we removed ATP from the reconstitution mix, we observed a significantly increased accumulation on the newly formed MT lattice compared to experiments with ATP (Fig.4E, F, Fig. S5C, D). Importantly, no accumulation on the minus-end was detected, and KIF2A clearly avoided binding to the GMPCPP-derived MT seed (Fig.4E, F, Fig. S5C, D). In none of our experiments did we observe accumulation of KIF2A at plus-ends (Fig. 4, Fig. S5, Movie S6). We conclude that in the presence of ATP KIF2A is an autonomous -TIP, in line with recent results (Henkin et al., 2023). However, in the absence of ATP KIF2A alters its binding mode, preferring the newly formed MT lattice over the MT minus-end, and avoiding binding to the seed. The latter data suggest that KIF2A prefers compacted GDP-over expanded GTP-MT lattices.

### Organisation and regulation of ICB MTs

The ICB is maintained for hours in mESCs, and ICB MTs are constantly moving inside a mESC colony (Movie S7). We therefore asked how ICB MTs are organised and regulated. To analyse total MT mass, we fixed mESCs and stained them with antibodies against alpha-tubulin. We used antigen retrieval to uncover the tubulin epitope in ICB MTs, since in fixed mESCs without antigen retrieval we hardly observed ICB MT staining, in contrast to the prominent staining of other MT-based structures (Fig. 1C). Despite antigen retrieval, we still did not detect tubulin signal inside the midbody of fixed mESCs (Fig. 5A, B), even though MTs are present, as shown using the SiR-Tubulin marker. Nevertheless, in cells lacking KIF2A the level of tubulin was clearly decreased in the arms of ICB MTs (Fig. 5A, B). These results suggest that there are less MTs in the ICB of KIF2A-depleted mESCs compared to wild-type cells, consistent with our live-imaging data using SiR-Tubulin. Taken together, our data strongly support the hypothesis that KIF2A stabilises ICB MTs.

**Figure 5.**
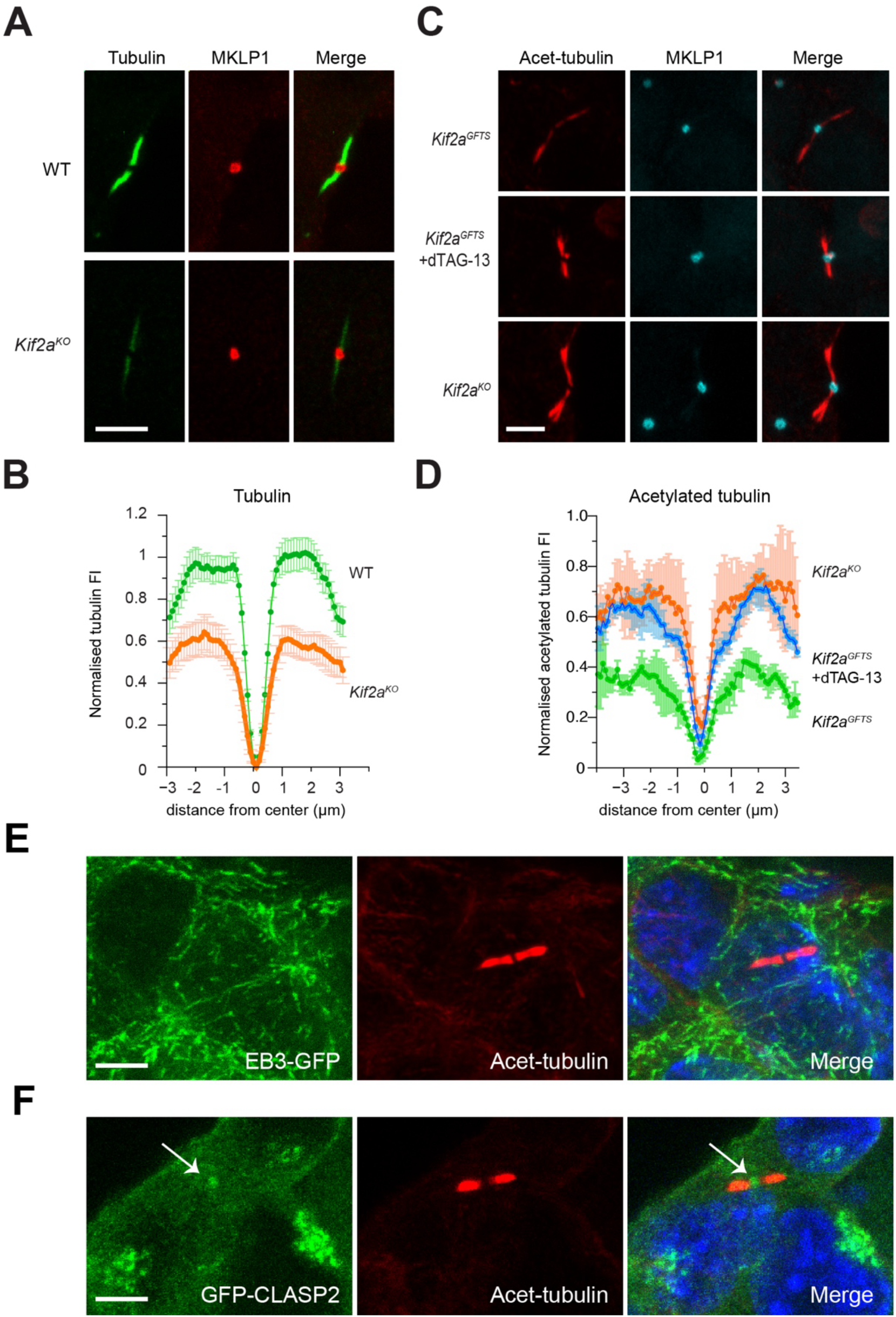
KIF2A regulates ICB MT behaviour. **A, B**) MT mass during cytokinesis in control and KIF2A-depleted mESCs. In (**A**) representative images are shown of ICB MTs stained with anti-alpha tubulin antibodies (green) and MKLP1 (red) in the indicated mESC lines, after antigen retrieval. Scale bar: 5 μm. In (**B**) line plots depict the averaged tubulin fluorescence over ICB MTs in wild type (WT, green), or *Kif2a^KO^* (orange) mESCs. ICBs were aligned based on MKLP1 signal (midbody marker). Data were normalised with min-max normalisation between 0 and 1. N = 2 independent experiments, n = 33 (KO) and 29 (WT). **C, D**) Acetylated tubulin distribution on ICB MTs. In (**C**) representative images are shown of ICBs stained with acetylated tubulin and MKLP1 in the indicated mESC lines. Scale bar: 5 μm. In (**D**) line plots depict the average acetylated tubulin fluorescence over the ICB in control (green), dTAG-13-treated (blue), or *Kif2a^KO^* (orange) mESCs. ICBs were aligned based on MKLP1 signal (midbody marker). Data were normalised with min-max normalisation between 0 and 1. Data were pooled from experiments with five mESC lines, i.e. control: wild type, *Kif2a^GFTS^* and *Kif2a^MFTS^* (mMaple3-tag instead of an eGFP-tag) mESCs, dTAG-13-treated: *Kif2a^GFTS^* and *Kif2a^MFTS^* mESCs treated with dTAG-13, *Kif2a^KO^*: two independent *Kif2a^KO^* lines. N = 3 independent experiments, n = 205 – 227 ICBs per condition. **E, F**) Localisation of selected +TIPs during cytokinesis. Representative images showing the localisation of EB3-GFP (**E**) or GFP-CLASP2 (**F**) in fixed mESCs, stained also with antibodies against acetylated tubulin. White arrows in (**F**) indicate GFP-CLASP2 signal in the midbody region. Scale bars = 3 μm.

Since ICB MTs are tightly packed bundles that are constantly moving and bending (Movie S7), we reasoned that they might undergo changes in interdimer distance, as well as mechanical, friction-induced damage, and we therefore tested the level of acetylated tubulin in ICB MTs. In stark contrast to total tubulin, we detected almost twice as much acetylated tubulin in KIF2A-depleted mESCs compared to control cells (Fig. 5C, D). These results suggest that the presence of KIF2A at ICB MTs prevents excessive ICB MT acetylation in mESCs. In line with this view, we often observed that the distribution of acetylated tubulin on the ICB arms was lower adjacent to the midbody (MKLP1 peak), where KIF2A-GFTS accumulated more strongly (Fig. 1D). Conversely, the acetylated tubulin signal was higher in distal regions of ICB MTs where KIF2A-GFTS accumulation was less pronounced (Fig. 1D). In conclusion, KIF2A maintains ICB MT mass and prevents excessive acetylation of ICB MTs. Given this surprising result we wondered how ICB MTs are organised and maintained.

To view dynamically growing MTs, we examined EB3, a prominent +TIP that marks the ends of growing MTs (Stepanova et al., 2003), and CLASP2, a MT growth-promoting +TIP (Yu et al., 2016). To avoid antibody penetration issues, we used mESC lines expressing EB3-GFP and GFP-CLASP2 for this analysis. Strikingly, we did not detect EB3-GFP signal on ICB MTs at all (Fig. 5E), whereas GFP-CLASP2 was only detected in the midbody region (Fig. 5F). These results suggest that ICB MTs are stable structures, and that their longitudinal extension during the first phase of cytokinesis occurs by EB-independent MT growth.

### KIF2A regulates pluripotency maintenance

mESCs can be maintained in a naïve, or ground state in defined medium containing the two inhibitors CHIR99021 and PD0325901 (referred to as 2i) and Leukaemia Inhibitory Factor (LIF) (Ying et al., 2008). As cytokinesis duration has been shown to be linked to pluripotency exit, (Chaigne et al., 2020) we performed a pluripotency exit and colony formation assay (Mulas et al., 2019) in control and KIF2A-depleted naïve mESCs. We found that *Kif2a^KO^* mESCs were approximately half as efficient in reverting to pluripotency after 24 hours of 2i/LIF removal (exit) compared to wild-type mESCs (Fig. 6A, B). This was also true when exit was slowed down by addition of IWP2, a Wnt inhibitor (ten Berge et al., 2011). These results suggest that KIF2A depletion causes premature exit from pluripotency.

**Figure 6.**
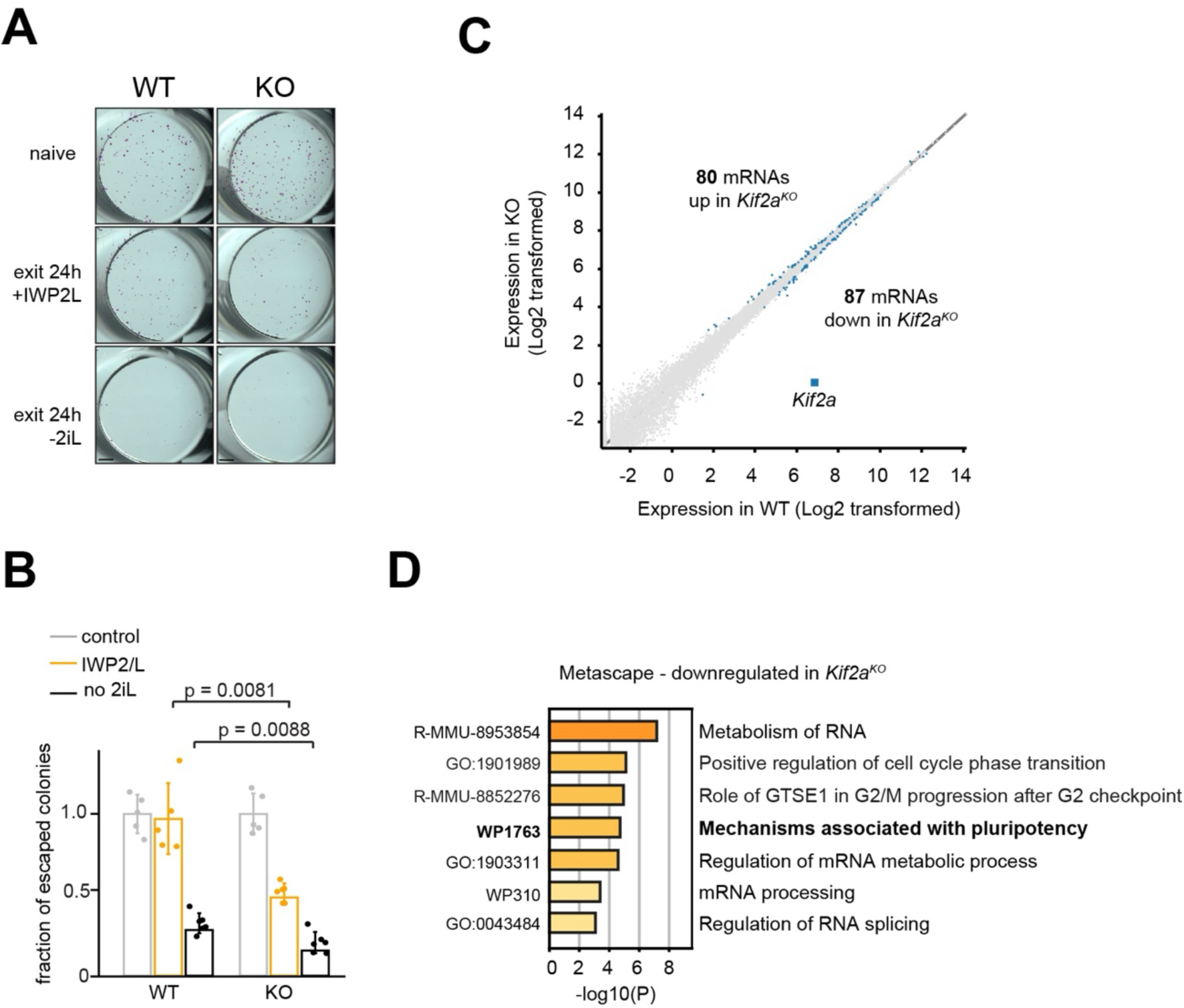
KIF2A regulates pluripotency exit. **A, B**) Pluripotency exit and colony formation assay. Wild type (WT) and *Kif2a^KO^*(KO) mESCs growing in naïve conditions, were trypsinised and replated in N2B27 exit medium (either lacking 2i/LIF or with IWP2 and LIF) for 24 hours followed by replating in naïve medium for 5 days. mESC colonies were then stained using alkaline phosphatase. In (**A**) representative alkaline phosphatase stainings are shown. The number of alkaline phosphatase-positive colonies was counted and plotted as a fraction of the number of colonies formed by naïve cells (set to 100%); this is shown in (**B**). Scale bar = 2 mm. P-values shown are derived from T-tests (n = 5 for controls and LIF +IWP2, n = 6 2i/LIF, 2 independent experiments). **C**) Differential gene expression analysis. We used DESeq2 to analyse differentially expressed genes (p<0.05) in WT versus KO mESCs. **D**) Metascape analysis of differentially expressed genes in WT versus KO mESCs. The 87 mRNAs that were found to be significantly downregulated in *Kif2a^KO^*(KO) mESCs were analysed by Metascape. Selected pathways are shown.

We next performed RNA-Sequencing (RNA-Seq) to uncover mechanisms underlying the mode(s) of action of KIF2A in an unbiased manner. For this, we compared RNA from two independently edited *Kif2a^KO^* lines (KO1 and KO2) and wild-type (WT) mESCs (see Table S2 for normalised RNA-Seq reads). Principal component analysis (PCA) showed segregation of triplicate samples by genotype (Fig. S6A). A differential gene expression analysis using DESeq2 yielded 80 upregulated and 87 downregulated mRNAs in KO mESCs as compared to WT (Fig. 6C). Metascape analysis (Zhou et al., 2019) of these differentially expressed genes (DEGs) revealed several pathways and terms enriched in KO mESCs, including “Mechanisms associated with pluripotency” (Fig. 6D). Thus, pluripotency-related mRNAs are significantly deregulated in *Kif2a^KO^* mESCs. Metascape analysis also uncovered pathways related to RNA biology (Fig. 6D), which is intriguing given the potential RNA-binding capacity of KIF2A in mESCs (Mallam et al., 2019).

Whereas DESeq2 examines differential gene expression at the level of individual mRNAs, Gene Set Enrichment Analysis (GSEA) determines whether a defined set of genes shows differences between two samples (Subramanian et al., 2005). We assembled a set of genes encoding proteins involved in centrosome regulation and MT nucleation, including all tubulin isotypes, the gamma-tubulin ring complex (γTuRC) and associating proteins (Bohler et al., 2021; Jackson, 2014; Yan et al., 2014), proteins involved in abscission (Chmp4b, Pdcd6ip, Cep55, Capn7, and Ist1 (Carlton et al., 2012; Chaigne et al., 2020; Paine et al., 2023)), as well as Cep170, an established KIF2A interaction partner (Zhang et al., 2019). We then used GSEA to assess whether KIF2A depletion affects mRNAs encoding these MT homeostasis and abscission factors. Interestingly, GSEA on this local set indeed revealed deviations in the two KO lines compared to wild-type, with several abscission factors being upregulated (Fig. S6B, Table S3). Combined, these data indicate that KIF2A depletion affects the pluripotent transcriptome.

## Discussion

Here, we combined cellular and *in vitro* approaches to gain insight into the function of the kinesin-13 member KIF2A in naïve mESCs. We find that in metaphase mESCs KIF2A localises near spindle poles, and our observations in KIF2A-depleted cells are consistent with KIF2A acting as a minus-end MT depolymerase at metaphase spindle poles. However, during cytokinesis, the dynamic behaviour of KIF2A changes, with KIF2A localising over the entire length of ICB MTs, suggesting strong binding to the lattice. FRAP results support the view of a differential binding mode of KIF2A near centrosomes (minus-end binding) compared to ICB MTs (tight lattice binding). Thus, KIF2A displays a dual binding behaviour in mESCs. This translates into a dual function, with KIF2A unexpectedly converting from a MT depolymerase in metaphase to a net MT stabiliser during cytokinesis. In this latter role KIF2A prolongs the duration of cytokinesis. Strikingly, increasing MT stabilisation by the taxol-derivative SiR-Tubulin also prolongs cytokinesis duration in wild type mESCs. Thus, two independent lines of evidence suggest that MT stabilisation at the ICB controls cytokinesis duration.

Our *in vitro* studies suggest that in the presence of ATP KIF2A is an autonomous -TIP. These results are similar to a recent report (Henkin et al., 2023). In addition, we find that in the absence of ATP KIF2A prefers the MT lattice over the minus-end. Thus, by inactivating KIF2A a switch in its MT binding mode is induced. We furthermore observed that KIF2A avoids the GMPCPP seed. Since GMPCPP is a GTP analog that maintains the MT lattice in an expanded state (Zhang et al., 2018), our data suggest that KIF2A binds the compacted MT lattice and not the expanded one. Altogether, our *in vitro* results indicate that the high affinity of KIF2A for ICB MTs might be due to inactive KIF2A binding to MTs with compacted lattices.

Using LSFM we were able to acquire high-resolution time lapse movies in mESC colonies over long periods of time. Imaging revealed that in naïve mESCs cytokinesis lasts 5-6 hours, which encompasses about one-third of their total cell cycle time (Soochit et al., 2021). We found that the ICB MT arms of future daughter cells display asymmetric bouts of KIF2A-GFTS and SiR-Tubulin, indicating that arms act independently. These asymmetric bouts might represent temporarily altered MT behaviour or lattice compaction state in one daughter cell in response to mechanical forces. Despite these bouts, our experiments strongly suggest that ICB MTs are stable, long-lived structures. For example, we did not detect EB3-related MT growth events along ICB MTs and no EB3-GFP was observed in the midbody. In addition, CLASP2 signal was restricted to the midbody, indicating that any MT growth promoted by CLASP2 occurs in the midbody and not elsewhere on ICB MTs. Thus, despite their long life time the ICB MTs of naïve mESCs are different from the long-lived MT bridges observed in the early embryo *in vivo*, in which dynamic MT plus ends were found to be present (Zenker et al., 2017).

LSFM imaging furthermore revealed two phases in cytokinesis, an initial one wherein ICB MT length increases, and a second phase, wherein length decreases. Concomitant with ICB MT length increase we observed an increase in KIF2A-GFTS accumulation on ICB MTs. In addition, we showed that ICB MTs are continuously moving, suggesting they undergo significant mechanical stress. Based on our FRAP results we propose that KIF2A binds the lattice of ICB MTs with high affinity and thereby maintains ICB MT stability. This is particularly important for the first phase of cytokinesis, as this phase is most affected in KIF2A-depleted mESCs. KIF2A might act simply by preventing the binding of factors that negatively regulate MT stability, for example severases. Alternatively, tight binding of KIF2A to ICB MT lattices in mESCs could help to maintain the compacted state. This behaviour is remarkably similar to that of Tau, a MAP with intrinsically disordered domains which was shown to form cohesive structures around neuronal MTs, thereby altering interdimer spacing (Siahaan et al., 2022). αTAT1 acts preferentially on MTs with an expanded MT lattice (Shen and Ori-McKenney, 2024), and the presence of more acetylated tubulin in ICB MTs of KIF2A-depleted mESCs could indicate that in the absence of KIF2A ICB MTs become more expanded and hence better substrates for αTAT1. The increased acetylation of ICB MTs in KIF2A-depleted mESCs might in turn serve as a back-up mechanism for MT protection (Portran et al., 2017). This would explain why the cytokinesis phenotype of KIF2A-depleted mESCs is relatively mild.

We have shown that in naïve mESCs KIF2A positively regulates ICB MTs, thereby prolonging cytokinesis and postponing abscission. Accordingly, KIF2A-depleted mESCs show premature pluripotency exit, coinciding with deregulated levels of mRNAs encoding pluripotency and other factors. Our data establish a link between MT stabilisation in cytokinesis and a pluripotent mESC transcriptome. Recently, the midbody was shown to be involved in local mRNA translation and to harbor mRNAs encoding proteins involved in cytokinesis, cell fate and pluripotency (Park et al., 2023). KIF23 was proposed to function in midbody RNA localisation by binding mRNAs with a disordered domain and coupling RNAs to MTs (Park et al., 2023). Interestingly, KIF2A also possesses disordered domains and has been shown to bind to RNA in mESCs (Mallam et al., 2019). KIF2A therefore, might be involved in localising mRNAs to ICB MTs. Future efforts will be directed at examining a possible RNA-related function of KIF2A.

## Materials & Methods

### Antibodies

We used the following primary antibodies in this study (dilutions and applications are also listed): mouse anti-acetylated tubulin (Sigma, T6793, immunofluorescent staining (IF) 1:1000), mouse anti-beta-tubulin (Sigma, T8328, western blot (WB) 1:1000), rabbit anti-beta-tubulin (Abcam, ab6046, IF 1:200), rat anti-alpha-tubulin (Abcam, ab6160, WB 1:4000, IF 1:1000), mouse anti-gamma-tubulin (Sigma, T6557, IF 1:300), rabbit anti-KIF2A (Novus Biologicals, NB500-180, WB 1:1000), rabbit anti-MKLP1 (Abcam, ab174304, IF 1:500), mouse anti-GAPDH (Absea, KT186, WB 1:2000). We used the following secondary antibodies: goat anti-mouse IgG Alexa Fluor 594 (Invitrogen, A11005), goat anti-rabbit IgG Alexa Fluor 647 (Invitrogen, A21244), goat anti-mouse IgG Alexa Fluor 647 (Invitrogen, A21236), goat anti-rabbit IgG Alexa Fluor 594(Invitrogen, A11012), IRDye 680RD goat anti-mouse IgG (LI-COR, 926-68070), IRDye 800RD goat anti-mouse IgG (LI-COR, 926-32210), IRDye 680RD goat anti-rabbit IgG (LI-COR, 926-68071), IRDye 800RD goat anti-rabbit IgG (LI-COR, 926-32211). Secondary Invitrogen antibodies were used at 1:1000 and secondary LI-COR antibodies were used at 1:15.000.

### Standard molecular biology methods

For RNA isolation, cells were collected and resuspended in 1 ml TRIzol (Sigma) and incubated for 5 min at 30 °C. 200 μl phenol-chloroform (Sigma) was added and samples were incubated on a shaker for 3 min at 30 °C. Samples were subsequently centrifuged for 15 min, 12.000 rpm at 4 °C in a table-top centrifuge. The aqueous phase (top layer) was collected and 250 μl 100% ethanol (Sigma) was added. Samples were then transferred to RNeasy spin columns (Qiagen) and processed according to the manufacturer’s instructions.

SDS-PAGE was carried out according to standard procedures using the mini-PROTEAN system (Biorad). After electrophoresis gels were either fixed and stained with Coomassie Brilliant Blue R-250 or blotted onto a PVDF (Millipore) membrane through wet-transfer for 2 hours at 4°C. The membrane was then blocked with 5% skim milk (Sigma) in phosphate-buffered saline (PBS) and 0.1% Tween-20 (PBS-T), for 30 minutes, and incubated overnight (O/N) at 4°C with primary antibody. Membranes were washed 3 times with PBS-T and incubated for 45 minutes with secondary antibody. After 3 washes with PBS-T, membranes were imaged using the Odyssey CLx (Licor).

### Cell culture and treatments

HEK293T cells were maintained in DMEM (Gibco) with 10% fetal bovine serum (FBS, Capricorn Scientific) and 1% penicillin/streptomycin (P/S, Sigma) at 37 °C, 5% CO_2_. HEK293T cells were transfected with X-tremeGENE HP DNA transfection agent (Roche) with a 1:2 DNA:reagent ratio and incubated for 24 – 48 hours before continuing with other experiments.

Experiments with mESCs were performed in specified media, which either contained two inhibitors (2i), i.e. 1 *μ*M PD0325901 (Stemgent, 04-0006) and 3 *μ*M CHIR99021 (Tocris, 4423-10), as well as serum and LIF, or just serum and LIF. For next generation sequencing experiments mESCs in 2i/L were used. For fluorescence microscopy experiments live mESCs were grown on 0.2% gelatin-coated plates or coverslips, or in chambers, in serum-free N2B27 medium (1:1 DMEM/F12-Glutamax (Gibco):Neurobasal (Gibco) medium, 1x N-2 supplement (Gibco, 17502001), 1x B27 minus vitamin A (Gibco, 12587010), 5 μM β-mercaptoethanol, 12.5 ng/ml Insulin (Sigma,19278), 1% P/S, 1000 U/ml LIF), supplemented with 2i. For IF and WB experiments, cells were plated on 0.2% gelatin-coated plates or coverslips in medium containing serum and LIF.

Transfection of mESCs for gene targeting experiments (see below) was performed with Lipofectamine 2000 (ThermoFisher, 11668030) with a DNA:reagent ratio of 1:4, at 37 °C, 5% CO_2_ on a shaker at 150 rpm to prevent cell attachment. After 15 minutes incubation cells were plated on iMEFs. To deplete KIF2A-GFTS from cells, mESCs were treated with a concentration of 50 nM dTAG-13 (Tocris, 6605), unless specified otherwise. The mESCs were treated with dTAG-13 for the indicated times.

For the pluripotency exit and colony formation assay (abbreviated as clonogenicity assay) 200,000 cells growing in naïve (N2B27 medium plus 2i/L) conditions, were plated on a 6-well plate in exit medium. This was either N2B27 medium without 2i/L, or N2B27 medium with LIF + 1 *μ*M IWP2 (Stem Cell Technologies, 72124), as indicated. After 24 hours of exit, cells were trypsinised into a single-cell suspension, and replated on 24-well plates in triplicate at 300 cells per well in complete N2B27 medium containing 2i/L. As a positive control, cells were maintained in naïve conditions without exit, and replated like the rest. After 5 days, cells were fixed for 2 minutes in 4% paraformaldehyde (pH 7.4), washed once with PBS, then stained using the alkaline phosphatase staining kit from Millipore (scr004) according to the manufacturer’s instructions. Then, cells were washed once with milliQ water, and air-dried in the dark overnight. The entire well was imaged using an Olympus dissecting microscope and all Alkaline phosphatase positive colonies were counted. The number of colonies formed by naïve cells was set to 100%.

### Generation of targeted and transgenic mESC lines

To generate knock-in mESC lines expressing various KIF2A fusion proteins or knockout mESCs in which a large part of the *Kif2a* gene was deleted, short guide RNAs (sgRNAs) were designed using the CHOPCHOP online tool (https://chopchop.cbu.uib.no/) to target the last exon of the *Kif2a* locus (knock-in lines and knockout line, 5’-ACGCCAACTTAGAGGGCTCG-3’) or the second exon of *Kif2a* (knockout line only, 5’-AGGCCGAATACACCAAGCAA-3’). sgRNAs were ligated into the pSpCas9(BB)-2A-Puro plasmid (PX459, Addgene #62988). The pUC8 vector (Sigma) was used as a backbone for cloning of homology arms (∼500 base pairs) and inserts. Human FKBP12 (324 base pairs) was ordered as a gBlock (IDT DNA), eGFP was amplified from a previously published plasmid (Yu et al., 2016). The F36V mutation of FKBP12 together with the TEV-linker-StrepII sequence were introduced by PCR. Gibson assembly was used to ligate and clone all sequences. All plasmids were sequence-verified.

mESCs were transfected with PX459 and pUC8 plasmids as described above. 24 hours after transfection cells were selected with 1 μg/ml puromycin (Sigma) for 2 days. After recovery of 3-4 days putative knock-in mESCs were sorted using flow cytometry and plated at low density on a 10 cm dish (putative knockout mESCs were not sorted but plated as such). Single colonies were picked and grown in 96-well plates. DNA was extracted from individual clones with lysis buffer (50mM Tris-HCl pH 8.0, 5mM EDTA pH 8.0, 0.5% SDS, 0.3 mg/mL Proteïnase K). PCR was performed with primers located outside the homology arms (sequences available upon request). After verification by PCR, we performed Sanger sequencing and western blot analysis to validate correct homologous recombination.

For generating GFP-CLASP2 knock-in mESCs, the sgRNA 5’-CGCAGAAGT ACTCGGCGC CG CGG – 3’, targeting the ATG of the alpha-isoform of *Clasp2* was used. Homology arms, corresponding to 700 bp upstream and downstream, respectively, of the sgRNA cut site, were amplified from genomic DNA of mESCs. The StrepII-TEV-GFP sequence was synthesized (Integrated DNA Technologies, IDT) and cloned into pUC8 vector together with left and right homology arms using Gibson Assemby (NEB). For targeting, 600,000 ES cells were reverse-transfected with 1 μg of guide RNA and 2 μg of homology template vector. Cells were then plated on 10 cm dishes with iMEFs in ES medium with serum and LIF. 24 hours post-transfection, medium was refreshed and Puromycin was added at 1 μg /ml for 48 hours. Cells were sorted after 5 days for GFP expression using FACSaria III system. After sorting, cells were seeded on iMEFs in 10-cm dishes at low density. After a week, single colonies were picked and maintained in 96-well plates, and screened by PCR and western blotting.

To generate a transgenic mESC line expressing EB3-GFP, the construct (Stepanova et al., 2003) was cloned downstream of a chicken beta-actin (CAG) promoter, linearized with Pvu1 (NEB), and transfected into mouse ESCs cultured under serum + LIF conditions. Transfection was performed in 6-cm dishes using 7.5 μg of linearized DNA and 15 μg of Fugene HD transfection reagent (Promega) according to the manufacturer’s instructions. 24 hours later, cells were expanded into 3X10 cm dishes, and Puromycin (1 μg /ml, Sigma) was added. Cells were kept under selection for 5-7 days before picking clones. Selected clones were expanded in medium containing serum and 2i/LIF and verified for EB3 expression using a confocal spinning disk microscope.

### Protein purification

KIF2A (2118 bp) was amplified from HEK293T cDNA, which was synthesized from HEK293T RNA using the SuperScript IV Transcriptase kit (Invitrogen,18090050). Primers used for the KIF2A amplification are available on request. eGFP was amplified from a previously published plasmid (Yu et al., 2016). The StrepII-TEV sequence (126 bp) was ordered as gBlock fragment from IDT DNA with overhangs for insertion into the pcDNA3 backbone and ligation to eGFP. The TEV sequence is followed by a linker sequence that translates to GGSGG. The complete StrepII-TEV-eGFP-KIF2A construct was cloned into pcDNA3 (Invitrogen) by Gibson Assembly (NEB). Constructs were verified after cloning by Sanger sequencing.

StrepII-TEV-eGFP-KIF2A was transfected in HEK293T as described above. After 48 hours cells were washed with cold PBS, collected and lysed with lysis buffer (50mM HEPES pH 7.4, 300mM NaCl, 0.5% Triton X-100 (Sigma), 1x cOmplete protease inhibitor cocktail (Roche)) for 20 minutes on ice. The lysate was incubated with Streptactin sepharose beads (Cytiva) for 1 hour at 4 °C and washed 3 times with lysis buffer without protease inhibitor at 4 °C. Beads were resuspended in elution buffer (50mM HEPES pH 7.4, 150mM NaCl, 1mM MgCl_2_, 1mM EGTA, 1mM DTT, 2.5mM d-Desthiobiotin (Merck), 0.05% Triton X-100) and rotated on a rotor at 4 °C for 2 hours. Eluate was collected and aliquoted in assay-sized aliquots, flash-frozen and stored at -80 °C. KIF2A concentration was determined with a Bovine serum albumin (BSA, Sigma) standard curve on a Coomassie R-250-stained SDS-PAGE gel.

### *In vitro* MT reconstitution assay

*In vitro* MT reconstitution assays were carried out according to previously published procedures (Bieling et al., 2007; Leslie and Galjart, 2013), with some modifications. Fluorescently labelled MT seeds were generated by mixing 62% porcine brain tubulin (Cytoskeleton, T240B), 16% X-rhodamine tubulin (Cytoskeleton, TL620M-A) or Hi-Lyte 647 Tubulin (Cytoskeleton, TL670M-A), and 22% biotin-tubulin (Cytoskeleton, T333PA) with 0.7 mM GMPCPP (Jena Biosciences, NU-405S). Mixtures were incubated at 37 °C for 40 minutes to allow polymerisation. The mixture was afterwards centrifuged for 10 minutes at 100.000xg in a Beckman tabletop airfuge, the supernatant was removed and the pellet was gently resuspended in 30 μl warm MRB80 buffer (80 mM PIPES-KOH pH 6.8, 1mM EGTA, 4mM MgCl_2_). MT seed samples were divided into 1 μl aliquots and flash-frozen. Oxygen Scavenger (OS) mix was prepared by dissolving 5 mg of catalase (Sigma, c9322), 10 mg of glucose oxidase (Sigma, G7141), and 15 mg of DTT in 500 μl of MRB80 buffer. The solution was then flash-frozen in 1 μl aliquots.

Chambers were prepared one day before the reconstitution assay. Glass slides (Epredia) and 18x18 mm coverslips (VWR) were treated O/N with 1M KOH in 100% ethanol. Slides and coverslips were washed 6 times and stored in autoclaved water (MilliQ). On the day of the assay glass slides and coverslips were dried with filtered air and a coverslip was mounted onto the glass slide with double-sided tape to create a flow-chamber with a width of 5 mm and a volume of 10 μl.

The flow-chamber was functionalized with 1 chamber volume of PLL-PEG-biotin (SuSoS) for 5 minutes. Excess was washed away with 5 chamber volumes of warm MRB80 buffer. Two chamber volumes of Neutravidin (Invitrogen, A2666, 1 mg/ml in PBS) were then added for 3 minutes and afterwards chambers were again washed with 5 chamber volumes of warm MRB80. Seeds were diluted in warm MRB80, flowed into the chamber and incubated for 5 minutes. Unbound seeds were washed away with 2 chamber volumes of warm MRB80, and next 3 chamber volumes of 5 mg/ml κ-casein (Sigma, C0406) dissolved in MRB80, were added to block aspecific binding. A protein mixture of 20 μl that was centrifuged for 8 minutes at 100.000xg to remove insoluble material was injected into the chamber and the flow-chamber was immediately imaged afterwards. The protein mixture contained 15 μM porcine brain tubulin, 1.25 μM X-rhodamine tubulin, 0.5 mg/ml κ-casein, 1.25 mM GTP (Cytoskeleton), 0.15% (v/v) methylcellulose (Sigma) and 75 mM KCl (Sigma). For experiments 0.5 μl of the OS mix described above, and 25 mM D-(+)-glucose solution in water (Sigma, G8270), were added to the reaction mix. ATP (Cytoskeleton, BSA04001) was added in a concentration of 0.5 mM.

MT *in vitro* reconstitution assays were imaged using a Nikon Ti-Eclipse inverted microscope with a total internal reflection fluorescence (TIRF) unit, equipped with a PLAN APO TIRF 100x/1.49 NA oil objective and a QuantEM512C 512x512 pixels 16bit camera (Photometrics). The chamber was heated to 30 °C with stage top incubator and objective heating (Tokai Hit). Dual colour imaging was performed with a DV2 Beamsplitter Green/Red (MAG Biosystems). The 491 nm and 561 nm lasers were used to image eGFP and X-Rhodamine, respectively. A single frame of the 647-Tubulin-labelled seed was acquired prior to the start of the timelapse with the 633 nm laser, also using the DV2 Beamsplitter. Stream acquisition was used with 500 ms exposure with the Metamorph Imaging Software.

### Immunofluorescence (IF) stainings

mESCs were grown on coverslips coated with 0.2% gelatin. dTAG-13 was added to the cells 24 hours before fixation, unless otherwise indicated. Cells were either fixed for 10 minutes with 100% methanol at -20 ^0^C, followed by 10 minutes with 4% paraformaldehyde (PFA, Sigma) at room temperature (RT), or for 10 minutes at RT with 4% PFA only, and then washed 3 times with PBS. Cells were then permeabilized with PBS containing 0.15% Triton X-100 (Sigma) for 10 minutes, washed with PBS, and incubated with blocking buffer (PBS, 0.1% Tween-20, 1% Bovine serum albumin (BSA, Sigma)) for 30-60 minutes on a shaker (100 rpm). Cells were subsequently incubated with the indicated antibodies in blocking buffer, either for 1 hour at RT, or O/N at 4 °C. Cells were washed 3 times, for 10 minutes each time, in wash buffer (PBS, 0.1% Tween-20) and incubated with secondary antibodies for 1 hour at RT. Cells were washed 3 times in wash buffer, once in 70% ethanol, once in 100% ethanol and finally air dried. Cells were mounted on glass slides with ProLong Gold antifade with DAPI (Invitrogen, P36931).

For antigen retrieval (alpha-tubulin IF), fixed coverslips were immersed in 10 mM Sodium Citrate buffer, ph 6.0 with 0.05% Tween-20, which had been heated to 95°C. Coverslips were left in this hot solution for 30-45 min until they had cooled to room temperature. Coverslips were washed twice in wash buffer, blocking solution was added, and the rest of the procedure was performed as described above.

### Standard fluorescence-based microscopy experiments

Standard confocal imaging was performed using the Leica SP8 AOBS microscope equipped with PLAN APO 63x/1.4 NA oil objective. DAPI, eGFP or Alexa488, Alexa 594, and Alexa 647, were imaged using a 405 nm, 488 nm, 561 nm or 633 nm laser, respectively. Detection was either by a Photo Multiplier Tube (PMT) or a Hybrid detector (HyD) and a scCMOS camera (Leica DFC9000 GTC DLS, 2048 x 2048 pixels). All images were acquired at a resolution of 1024 X1024 pixels.

For live-cell imaging of mESCs from metaphase-to-telophase, mESCs were plated two days before the start of the experiment on µ-Dish 35 mm dishes (Ibidi, 81156), coated with 0.2% gelatin. The day before imaging the cells were incubated with dTAG-13 when necessary. Cells were incubated with SiR-tubulin dye as described below and kept at 37 °C, 5% CO_2_. SiR-tubulin was imaged using the 633 nm laser and KIF2A-GFTS with 488 nm. Z-stacks were acquired every 3 minutes with a 2 μm step-size.

Fluorescence recovery after photobleaching (FRAP) experiments were largely performed as described previously (Soochit et al., 2021), using a Nikon Ti-Eclipse inverted microscope with a CSU-X1 (Yokogawa) spinning disc equipped with a PLAN APO TIRF 100x/1.49 NA oil objective and a QuantEM512C 512x512 pixels 16bit camera (Photometrics). *Kif2a^GFTS^* mESCs were plated on 0.2% gelatin-coated coverslips the day before the FRAP experiment. Cells were heated to 37 °C with a stage top incubator and objective heating (Tokai Hit). Photo bleaching was performed using the iLAS FRAP module (Roper Scientific) integrated into Metamorph, with the 491 nm laser set at 100%. Region of interests (ROIs) were selected and bleached for 90ms with 10 repetitions. Subsequent images were acquired with 35% laser power for one minute, 2 frames per second with a 400 ms exposure. Eleven images were acquired before bleaching.

### Light sheet fluorescence microscopy (LSFM)

For long-term imaging 20.000 wild-type, *Kif2a^GFTS^* or *Kif2a^KO^* mESCs were plated 1 day before imaging on a TruLive3D dish (Ibidi, Bruker) that was plasma cleaned for 3 minutes with the Glow Plasma System (Glow Research) and coated with 0.2% gelatin for 15 minutes. On the day of the experiment, to visualise the MT network, 20 nM SiR-tubulin (Spirochrome) was added to the medium six hours prior to imaging. Afterwards, cells were washed once with PBS and regular medium without dye was added. The TruLive3D dish was inserted into the InVi SPIM sample holder and placed into the imaging chamber. The environmental control was set at 37 °C, 5% CO_2_ and 87% humidity, and lid heating was enabled. dTAG-13 treatment was performed for 24 hours.

The InVi SPIM fluorescence light sheet microscope (Luxendo, Bruker) is equipped with a Nikon CFI Plan Fluor 10x/0.3 NA illumination water objective and a Nikon CFI Apo 25x/1.1 NA detection water objective. For imaging of KIF2A-GFTS the 488 nm laser was set at 8% and combined with the 497-554 band pass filter. SiR-tubulin was imaged with 10-15% 633 nm laser and the 656 long pass filter. Beams were split with 560 nm dichroic switch. 3D images were acquired with the bessel light sheet with a thickness of 40, a line mode of 200 pixels and a magnification of 62.5x. Images were acquired every 10 min for a total of 16 hours with a 90 ms exposure, 10 ms delay and a z-step size of 1 μm with 2x high-speed sCMOS cameras (Hamamatsu Orca Flash 4.0 V3 2048 × 2048 pixels). For the *Kif2a^GFTS^* cells, all samples were imaged with the 488 and 633 nm laser, even when KIF2A-GFTS was depleted by dTAG-13 treatment. This way we could verify KIF2A-GFTS depletion in the dTAG-13-treated cells and keep experiments comparable.

### Image analysis

Standard FRAP analysis was performed either using the EasyFRAP-web application (https://easyfrap.vmnet.upatras.gr/) (Koulouras et al., 2018) or a simplified protocol. For the EasyFRAP-web application in each experiment the fluorescence intensity over time for three ROIs was determined, i.e. the ROI that was bleached (ICB or metaphase spindle pole), an area inside the same cell but outside the bleached ROI and an area outside the cell. For the simplified protocol we selected ICB structures of similar length and determined fluorescence intensity over time using Fiji (Schindelin et al., 2012). We examined three ROIs, i.e. the ROI that was bleached, a ROI on the non-bleached arm of the same ICB MT structure and a background ROI. We deducted background fluorescence intensity from the other ROIs. In both approaches a full-scale normalisation was performed to normalise for differences in starting intensity, total fluorescence, and bleaching depth. For the line analysis we drew a line with a pixel width of 10 over the complete ICB MT network and determined fluorescence intensity over time for the line.

Quantification of fluorescence intensities of immunofluorescence stainings was performed on unprocessed maximum intensity projection images. Intensity profiles of KIF2A, MKLP1, alpha-tubulin or acetylated tubulin at the ICB were determined by line plots tracing the ICBs manually. The intensity profiles were subsequently aligned based on the midbody, which was either a local minimum or the MKLP1 local maximum. Data for each experiment was normalised with a min-max normalisation between 0 and 1.

Metaphase-to-telophase duration in live mESCs was calculated based on mitotic MT structures visualized by SiR-tubulin. Time was measured from the start of metaphase, i.e. formation of the metaphase spindle, until the end of telophase, i.e formation of ICB MTs. Distance between spindle poles in metaphase was determined manually in Fiji with line plots.

For LSFM image analysis acquired data were visualised by opening stacks and cropping movies in x, y, z and time (t) using the BigDataProcessor plugin (Tischer et al., 2021). When necessary, channels were aligned by manual transformation in x and/or y. The surface area of colonies was determined by taking the image nearest the surface of the TruLive3D dish, and outlining the colony edge using fluorescent signal and the segmented line command in Fiji. Duration of cytokinesis was manually determined in maximum intensity projections of 3D movies. We either traced KIF2A_GFTS or SiR-tubulin signal, from the formation of ICB MTs until the disappearance of signal (which we defined as the end of cytokinesis), by following the ICB MTs in time.

For detailed analysis of KIF2A-GFTS and SiR-Tubulin dynamics, a subset of 10 cropped time lapse movies was selected in dTAG-13-treated and non-treated mESCs (i.e. 20 images in total). Lines were manually annotated in each frame time based on the SiR-tubulin channel, allowing us to extract the length and line intensity profile per time frame. To account for the thickness of the line profiles, 17 pixels in width were averaged (the annotated pixel plus 8 pixels on each side). Background intensity correction was applied by subtracting from the mean background intensity value per time frame.

### Mass spectrometry

SDS-PAGE gel lanes were cut into 2-mm slices and subjected to in-gel reduction with dithiothreitol, alkylation with iodoacetamide and digested with trypsin (sequencing grade; Promega), as described previously (Schwertman et al., 2012). Nanoflow liquid chromatography tandem mass spectrometry (nLC-MS/MS) was performed on an EASY-nLC coupled to an Orbitrap Fusion Lumos Tribid mass spectrometer (ThermoFisher), operating in positive mode. Peptides were separated on a ReproSil-C18 reversed-phase column (Dr Maisch; 15 cm × 50 μm) using a linear gradient of 0– 80% acetonitrile (in 0.1% formic acid) during 90 min at a rate of 200 nl/min. The elution was directly sprayed into the electrospray ionization (ESI) source of the mass spectrometer. Spectra were acquired in continuum mode; fragmentation of the peptides was performed in data-dependent mode by HCD.

Raw mass spectrometry data were analyzed with the MaxQuant software suite ((Cox et al., 2009); version 2.1.3.0) as described previously (Schwertman et al., 2012) with the additional options ‘LFQ’ and ‘iBAQ’ selected. The false discovery rate of 0.01 for proteins and peptides and a minimum peptide length of 7 amino acids were set. The Andromeda search engine was used to search the MS/MS spectra against the Uniprot database (taxonomy: *Homo sapiens*, release May 2022) concatenated with the reversed versions of all sequences. A maximum of two missed cleavages was allowed. The peptide tolerance was set to 10 ppm and the fragment ion tolerance was set to 0.6 Da for HCD spectra. The enzyme specificity was set to trypsin and cysteine carbamidomethylation was set as a fixed modification. Both the PSM and protein FDR were set to 0.01. In case the identified peptides of two proteins were the same or the identified peptides of one protein included all peptides of another protein, these proteins were combined by MaxQuant and reported as one protein group. Before further statistical analysis, known contaminants and reverse hits were removed.

### RNA-sequencing

To prepare RNA samples we plated *Kif2a^KO^* and WT mESC lines on 0.2% gelatin-coated 6 cm plates two days before RNA preparation. Cells were then collected and RNA was isolated as described under ‘Standard molecular biology methods’. Total RNA was checked on an Agilent 2100 Bioanalyzer using a RNA nano assay. All samples had a RIN value higher than 9.2. RNA-Seq libraries were prepared according to the Illumina TruSeq Stranded mRNA Library Prep kit and sequenced according to the Illumina TruSeq Rapid v2 protocol on an Illumina NextSeq2000 sequencer, generating paired-end clusters of 50 bases in length. Illumina adapter and poly-A sequences were removed from the reads and the trimmed reads were then aligned to the GRCm38 mouse genome reference using HISAT2 (Kim et al., 2019).

Downstream analysis was performed using SeqMonk (https://www.bioinformatics.babraham.ac.uk/projects/seqmonk), followed by DESeq2 (Love et al., 2014), Metascape (Zhou et al., 2019), principal component analysis (performed with the 300 most variant genes) using Pcaexplorer (Marini and Binder, 2019), or Gene Set Enrichment Analysis (Mootha et al., 2003; Subramanian et al., 2005). We assembled a local GSEA dataset, consisting of a set of mRNAs encoding factors involved in centrosome regulation and MT nucleation, including the gamma-tubulin ring complex (γTuRC) and associating proteins (Bohler et al., 2021; Jackson, 2014; Yan et al., 2014), proteins involved in abscission (Chmp4b, Pdcd6ip, Cep55, Capn7, and Ist1 (Carlton et al., 2012; Chaigne et al., 2020; Paine et al., 2023)), all tubulin isotypes, and Cep170, an established KIF2A interaction partner (Zhang et al., 2019).

### Statistics and reproducibility

Independent student t-tests (also known as two-sample t-test) were performed when comparing the means of two samples. Analysis of statistical variance (ANOVA) was carried out when comparing multiple groups, which was followed either by a Tukey honest significant difference (HSD) post hoc test, or a Dunnett’s post hoc test in case of comparison of all groups to one control group. Statistical tests were performed in R. Repeated measures ANOVA was performed in Prism using the timepoints without missing values (GraphPad, Dotmatics). Each experiment (N) was carried out at least twice (exact N and sample sizes (n) are found in the figure legends). P-values are shown in the figures.

## Acknowledgements

This work was supported by grants from the Netherlands Organisation for Scientific Research (ZonMW TOP 40-00812-98-17045). H.K and I.S were funded by the Dutch Research Council (NWO) through the Building Blocks of Life research program (GENOMETRACK project, Grant No. 737.016.014). We thank Jan Sakoltchik for making the GFP-CLASP2 knock-in mESC line and Emiel van Genderen for his advice on the pluripotency exit experiment.

## Author Contributions

L.S. and S.B. performed the *in vitro* reconstitution experiments, the cellular experiments and analysis. B.v.H. generated the *Kif2a^KO^* mESC lines. L.S., H.K., I.S. S.B. and N.G. analysed the light-sheet data. L.S., J.D., D.D. and N.G. performed and analysed the mass spectrometry experiments. L.S., M.v.d.H., W.v.IJ. and N.G. performed and analysed the RNA-seq experiments. G-J.K. and B.D. helped in the light-sheet imaging experiments. L.S., D.H., S.B., and N.G. conceived experiments presented in this work. All authors contributed to the writing of this manuscript.

