## Supplemental Information for "KIF2A maintains cytokinesis in mouse embryonic stem cells by stabilising intercellular bridge microtubules"

**Supplementary Information contains:**

Supplementary Figures & Legends,

Supplementary Tables Legends,

Supplementary Movies Legends.

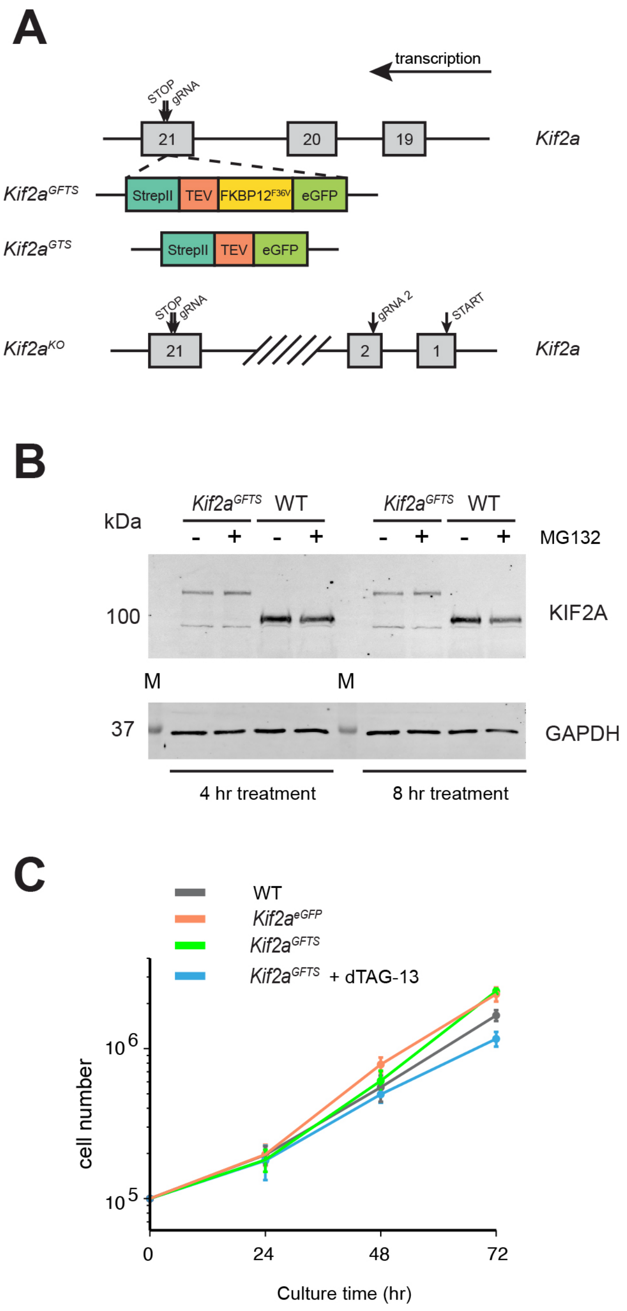
A
transcription
STOP
gRNA
21
20
19
Kif2a
Kif2a^GFTS
StrepII
TEV
FKBP12^F36V
eGFP
Kif2a^GTS
StrepII
TEV
eGFP
STOP
gRNA
gRNA 2
START
Kif2a^KO
21
2
1
Kif2a
B
kDa
Kif2a^GFTS
WT
Kif2a^GFTS
WT
- + - + - + - +
MG132
100
KIF2A
M
M
37
GAPDH
4 hr treatment
8 hr treatment
C
WT
Kif2a^eGFP
Kif2a^GFTS
Kif2a^GFTS + dTAG-13
cell number
10^6
10^5
0
24
48
72
Culture time (hr)

*Figure S1. Features of engineered mESCs.*

**A**) Schematic representation of CRISPR-cas9 gene-editing in mESCs. The *Kif2a*$^{GFTS}$ construct contains the following casettes (from 5' to 3'): eGFP, FKBP12$^{F36V}$, TEV (TEV protease cleavage site) and StrepII (Strep-tag II). The *Kif2a*$^{GTS}$ construct lacks the FKBP12$^{F36V}$ module. Constructs were targeted to exon 21 of the *Kif2a* locus, just before the STOP codon and were inserted in the order GFTS, or GTS in the *Kif2a* gene. The *Kif2a*$^{KO}$ cell line was generated with 2 sgRNAs targeting regions near the 5'end and 3'end of the *Kif2a* locus, resulting in the complete removal of exons 3-20, and partial removal of exons 2 and 21. **B**) Effect of proteasome inhibition on KIF2A levels. The indicated mESC lines (WT: wild type) were grown for 4 or 8 hr in the presence (+) or absence (-) of the proteasome inhibitor MG132 (1 μM). Cells were then lysed and KIF2A (in WT mESCs) or KIF2A-GFTS (in *Kif2a*$^{GFTS}$ mESCs) levels were analysed by western blot using anti-KIF2A (upper panel) or anti-GAPDH (lower panel) antibodies. Marker proteins (M) and relative molecular mass (in kDa) are shown to the left. **C**) Cell proliferation assay. The indicated mESC lines were plated (100,000 mESCs at onset of experiment) and cells were counted after 24, 48 and 72 hr. Shown are average cell numbers with SEM.

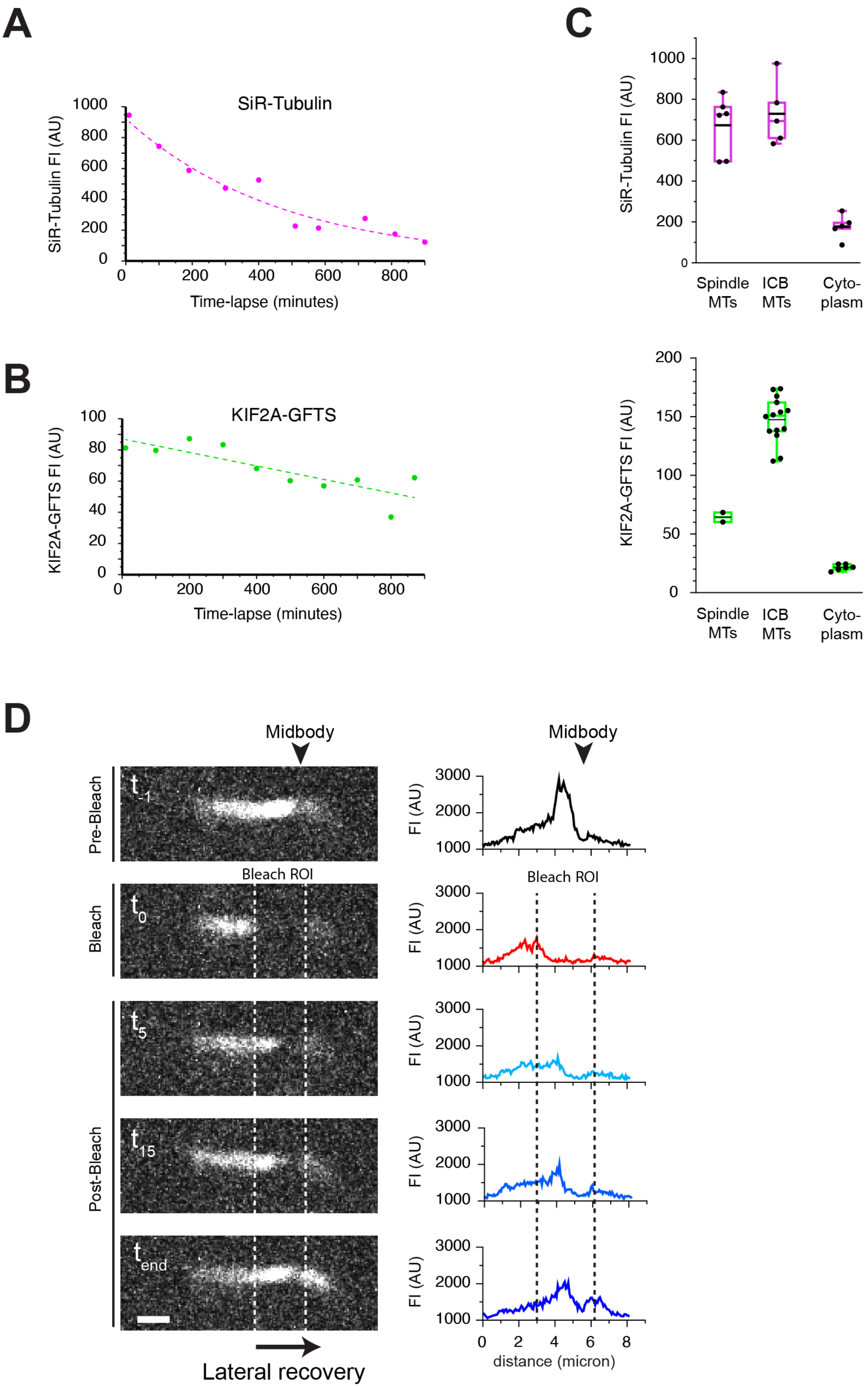

A
SiR-Tubulin
SiR-Tubulin FI (AU)
Time-lapse (minutes)
B
KIF2A-GFTS
KIF2A-GFTS FI (AU)
Time-lapse (minutes)
C
SiR-Tubulin FI (AU)
Spindle MTs
ICB MTs
Cyto-plasm
KIF2A-GFTS FI (AU)
D
Midbody
Pre-Bleach
Bleach
Post-Bleach
Bleach ROI
FI (AU)
Lateral recovery
distance (micron)

*Figure S2. Time-dependent decay of fluorescence intensities.*

**A, B**) SiR-tubulin (**A**) and KIF2A-GFTS (**B**) fluorescence intensity (FI) was measured in arbitrary units (AU) at selected time points during a complete 16 hr LSFM time lapse experiment. SiR-Tubulin FI was measured by placing a circular region of interest (ROI) over a centrosomal area at the onset of metaphase, when the SiR-Tubulin signal should be very high. KIF2A-GFTS signal was measured by placing a ROI over ICB MTs at the moment when KIF2A was most abundant. **C**) SiR-tubulin (upper panel) and KIF2A-GFTS (lower panel) fluorescence intensity (FI) was measured in arbitrary units (AU) at spindle and ICB MTs, and in the cytoplasm of mESCs, at the beginning of a LSFM time lapse experiment. Whisker plot shows interquartile range, median (purple/green lines), and average (black lines). **D**) Dynamic behaviour of KIF2A-GFTS at an individual ICB MT network. A line with pixel width 10 was drawn over an ICB MT network and the fluorescence intensity (FI) in arbitrary units (AU) of the whole ICB MT structure was measured over time. Depicted to the left are still images of selected time points, the stippled white lines indicate the bleach ROI borders. To the right plots are shown of the FI over the line of the corresponding still image. the stippled black lines indicate the bleach ROI borders. Note the lateral diffusion (from left to right) in time of KIF2A-GFTS over the ICB MTs. The position of the midbody is indicated with black arrowheads.

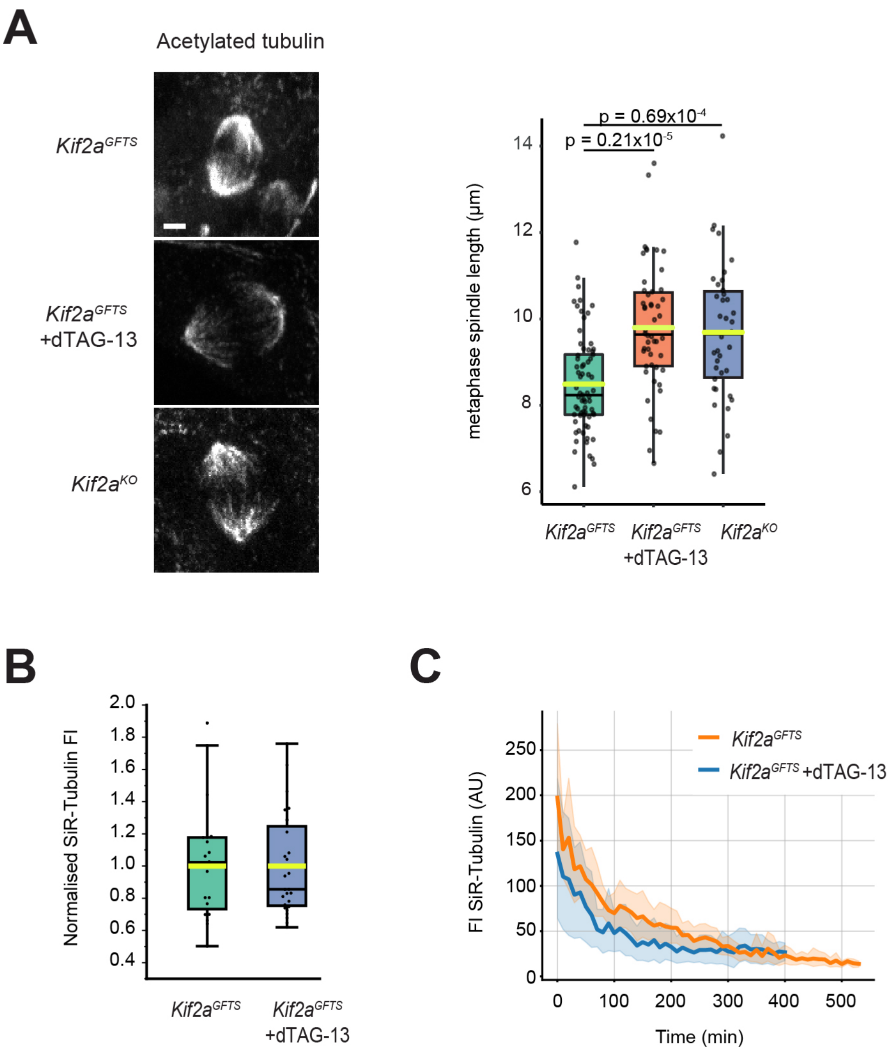


*Figure S3. Effects of KIF2A depletion on mitotic parameters.*

**A**) mESC metaphase phenotype after depletion of KIF2A-GFTS. Left panel: representative immunofluorescence image of acetylated tubulin pattern in in *Kif2a*$^{GFTS}$, dTAG-13-treated *Kif2a*$^{GFTS}$, and *Kif2a*$^{KO}$ mESCs (Scale bar = 2 µm). Right panel: whisker plot showing interquartile range, median (black lines), and average (yellow lines), and depicting the pole-to-pole metaphase spindle length in *Kif2a*$^{GFTS}$, dTAG-13-treated *Kif2a*$^{GFTS}$ and *Kif2a*$^{KO}$ mESCs. N = 3, n = 38-69 cells per condition. **B**) SiR-Tubulin accumulation on ICB MTs at the onset of cytokinesis. The fluorescence intensity (FI) of SiR-Tubulin was measured in the indicated mESC lines at the onset of cytokinesis (corresponding to t20 in Fig. 4a). Average FI values were normalised. Whisker plot shows interquartile range, median (black lines), and average (yellow lines). N= 20 (not-treated *Kif2a*$^{GFTS}$ mESCs), 27 (*Kif2a*$^{GFTS}$ mESCs treated with dTAG13). Data were obtained from 6-7 mESC colonies in 2 independent experiments. **C**) SiR-Tubulin behaviour on ICB MTs during cytokinesis. The fluorescence intensity (FI) of SiR-Tubulin was measured in arbitrary units (AU) in the indicated mESC lines during a complete imaging experiment. Lines depict average of all data points, shaded areas represent the SEM. n = 10 ICBs per condition.

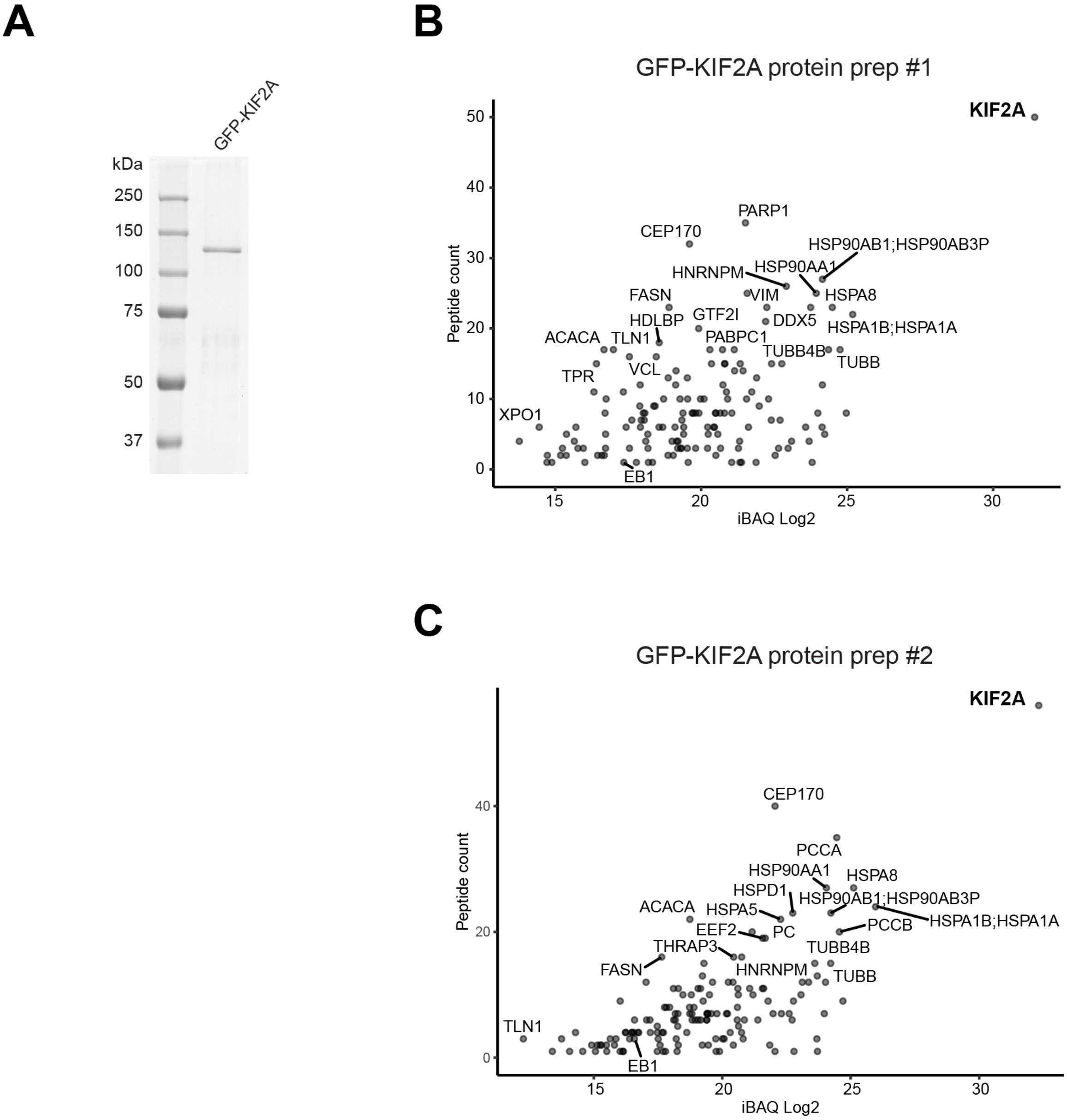


*Figure S4. In vitro behaviour of KIF2A*

**A**) Preparation of GFP-KIF2A. StrepII-tagged eGFP-KIF2A was produced in HEK293T cells and purified using Streptactin Sepharose beads. Shown is a Coomassie-stained SDS-PAGE gel loaded with purified GFP-KIF2A. Marker proteins with their relative molecular mass (in kDa) are shown to the left. **B, C**) Mass spectrometric-based analysis of purified GFP-KIF2A. StrepII-TEV-eGFP-KIF2A was independently purified two times (prep #1 (**B**), prep #2 (**C**)) from HEK293T cells and analysed by mas spectrometry. For a complete list of proteins, see Table S1. Based on iBAQ score GFP-KIF2A is at least 100-fold in excess in both preparations compared to co-purifying proteins.

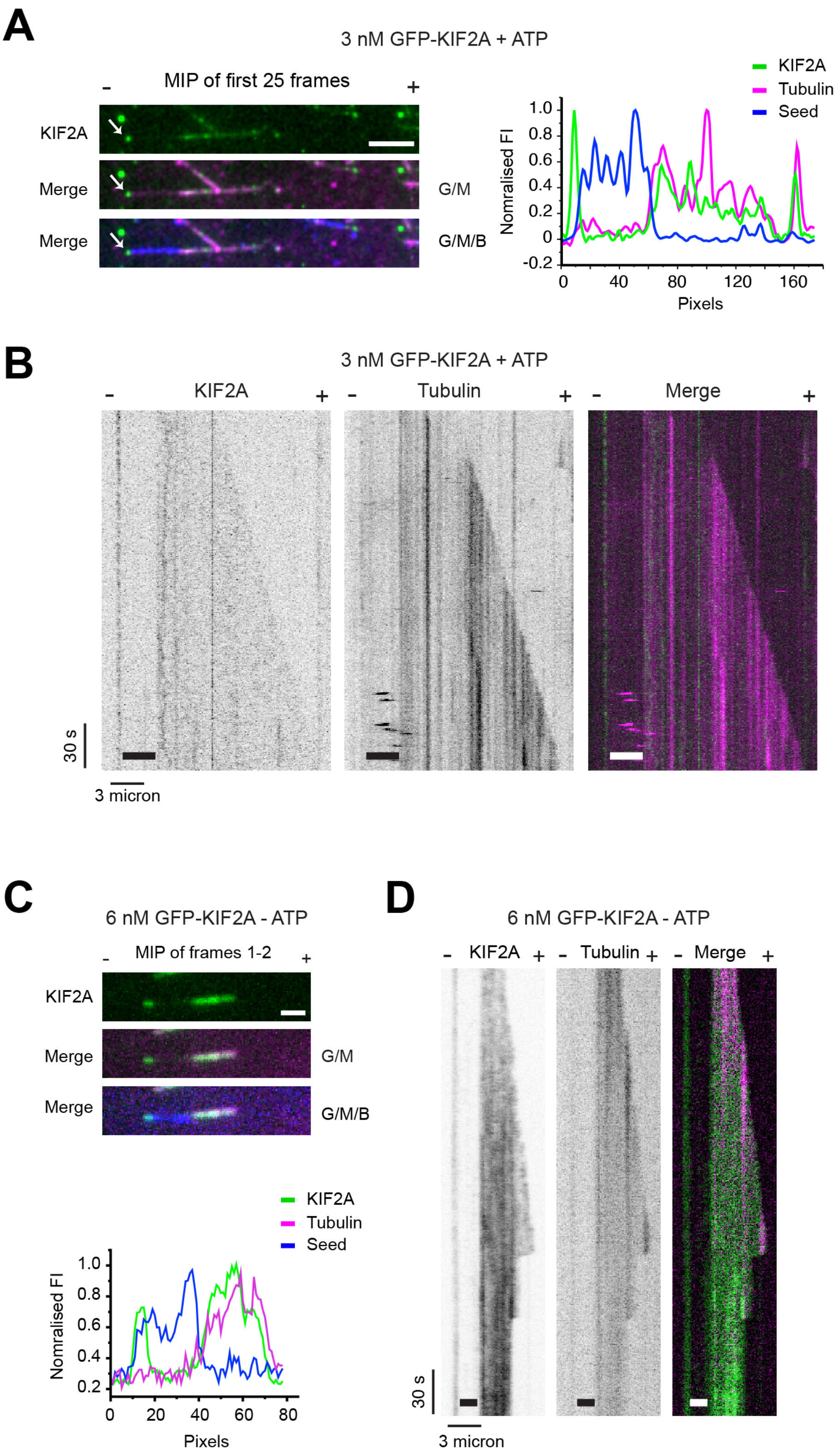
A
3 nM GFP-KIF2A + ATP
MIP of first 25 frames
KIF2A
Merge
Merge
G/M
G/M/B
KIF2A
Tubulin
Seed
Nomralised FI
Pixels
B
3 nM GFP-KIF2A + ATP
KIF2A
Tubulin
Merge
30 s
3 micron
C
6 nM GFP-KIF2A - ATP
MIP of frames 1-2
KIF2A
Merge
Merge
G/M
G/M/B
KIF2A
Tubulin
Seed
Nomralised FI
Pixels
D
6 nM GFP-KIF2A - ATP
KIF2A
Tubulin
Merge
30 s
3 micron

*Figure S5. GFP-KIF2A has affinity for MT minus-ends and the MT lattice.*
**A**) Maximum intensity projection (MIP) of TIRF microscopy images (first 25 frames) of 3 nM GFP-KIF2A binding to MTs. The experiment was carried out in the presence of ATP. GFP-KIF2A is labeled in green (G) and tubulin in magenta (M). The MT seed is labelled in blue (B). Note that the seed signal was captured in the first frame of the time lapse, and the still image was then superimposed. The - and + signs indicate MT minus- and plus-end, respectively. GFP-KIF2A accumulation on the MT minus-end is indicated by the white arrow. Normalised fluorescence intensity distributions (FI) of three dyes along the MT are plotted besides the MIP. Scale bar: 6 µm. **B**) Kymograph of *in vitro* MT reconstitution assay with 3 nM GFP-KIF2A. The experiment was carried out in the presence of ATP. In the merge, GFP-KIF2A is labeled in green and tubulin in magenta. The MT seed is indicated in the kymographs by a thick line. The - and + signs indicate MT minus- and plus-end, respectively. **C**) Maximum intensity projection (MIP) of TIRF microscopy images (frames 1 and 2) of 6 nM GFP-KIF2A binding to MTs. The experiment was carried out in the absence of ATP. GFP-KIF2A is labeled in green (G) and tubulin in magenta (M). The MT seed is labelled in blue (B). The - and + signs indicate MT minus- and plus-end, respectively. The seed signal was captured in the first frame of the time lapse, and the still image was then superimposed. Normalised fluorescence intensity distributions (FI) of the three dyes along the MT are plotted below the MIP. Comparison of these normalised FI distributions with those obtained in the presence of ATP (panel A, see also Figure 4A, C) shows that in the absence of ATP GFP-KIF2A does not accumulate at the very distal minus-end but instead binds the lattice. Scale bar: 2 µm. **D**) Kymograph of *in vitro* MT reconstitution assay with 6 nM GFP-KIF2A. The experiment was carried out in the absence of ATP. In the merge, GFP-KIF2A is labeled in green and tubulin in magenta. The MT seed is indicated in the kymographs by a thick line. The - and + signs indicate MT minus- and plus-end, respectively.

A

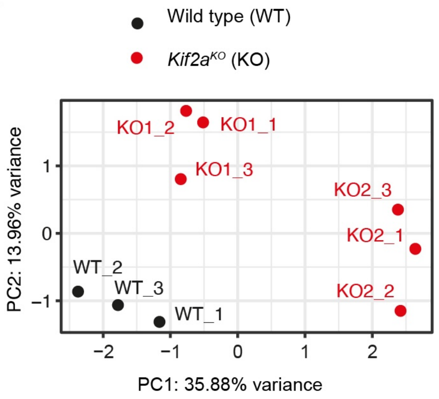
Wild type (WT)
Kif2a^KO (KO)
KO1_2
KO1_1
KO1_3
KO2_3
KO2_1
KO2_2
WT_2
WT_3
WT_1
PC2: 13.96% variance
PC1: 35.88% variance
1
0
−1
−2
−1
0
1
2

B

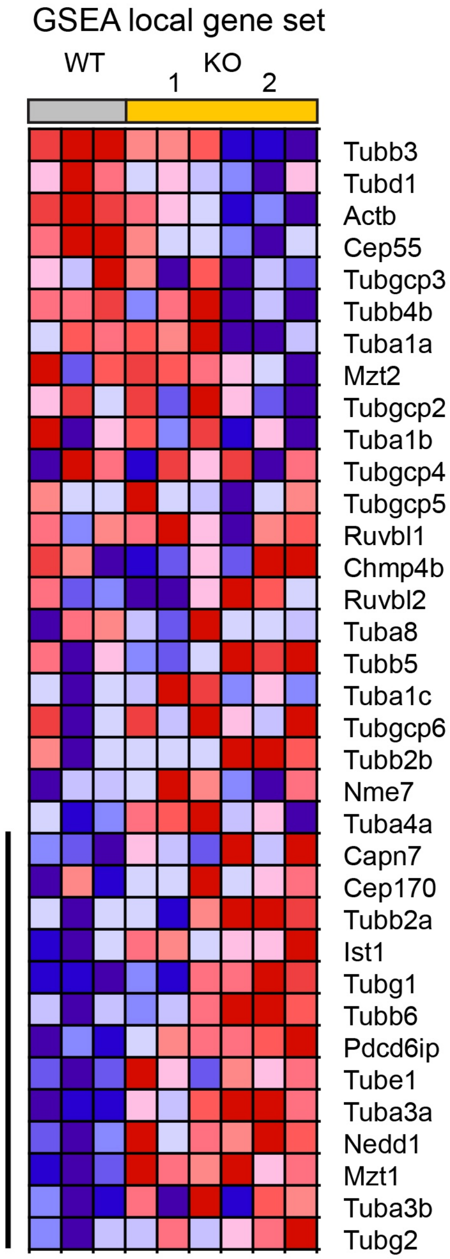
GSEA local gene set
WT
KO
1
2
Tubb3
Tubd1
Actb
Cep55
Tubgcp3
Tubb4b
Tuba1a
Mzt2
Tubgcp2
Tuba1b
Tubgcp4
Tubgcp5
Ruvbl1
Chmp4b
Ruvbl2
Tuba8
Tubb5
Tuba1c
Tubgcp6
Tubb2b
Nme7
Tuba4a
Capn7
Cep170
Tubb2a
Ist1
Tubg1
Tubb6
Pdcd6ip
Tube1
Tuba3a
Nedd1
Mzt1
Tuba3b
Tubg2

*Figure S6. Transcriptomic analysis of control and KIF2A-depleted mESCs.*

**A**) Principal component analysis of RNA-Seq datasets. Principal component analysis (PCA) was performed on the top-300 deregulated genes. We compared RNA derived from wild type (WT) and two independent *Kif2a*$^{KO}$ (KO) mESC lines. We analysed three biological replicates per line. **B**) Supervised GSEA analysis of KIF2A function in mESCs. Control and KIF2A-depleted RNA-Seq samples were subjected to a supervised GSEA where we locally assembled mRNAs encoding a set of factors involved in centrosome regulation, MT nucleation, and abscission, as well as a KIF2A-interacting partner, and asked whether mRNAs in this set were deregulated. We observed deviation of mRNA levels from control, as shown by the heatmap (red: high expression, blue: low expression). The black line indicates genes significantly contributing to the GSEA score.

*Table S1 - KIF2A fusion protein mass spectrometry analysis.*

StrepII-TEV-eGFP-KIF2A was independently purified two times (prep #1, prep #2) from HEK293T cells and analysed by mass spectrometry. All proteins with a Mascot score less than 20 were removed, as were obvious contaminants such as keratins. Displayed are IBAQ Log2 scores and number of peptides detected.

*Table S2 - RNA-Sequencing.*

mRNA was isolated from a wild type (WT) mESC line and from two independently gene-edited mESC *Kif2a*$^{KO}$ (KO) lines. RNA-Seq was performed on triplicate samples, shown are RPKM values.

*Table S3 - GSEA Local Dataset.*

We locally assembled mRNAs encoding a set of factors involved in centrosome regulation, MT nucleation, and abscission, as well as a KIF2A-interacting partner. Using GSEA we asked whether mRNAs in this set were deregulated. Shown are the rank of the genes in a supervised GSEA, as well as rank metric scores, running enrichment score (ES) and core enrichment. See Fig. S6B for corresponding heatmap.

*Movie S1*

z-stack (35 slices, 1 micron per slice) of a mESC colony expressing KIF2A-GFTS (green) and incubated with SiR-tubulin (red). ICB MTs are present throughout the colony.

*Movie S2*

3D projection of the same mESC colony shown in Movie S1 expressing KIF2A-GFTS (green) and incubated with SiR-tubulin (red).

*Movie S3*

LSFM-based time lapse imaging of *Kif2a*$^{GFTS}$ mESCs incubated with SiR-tubulin. The mESC colony was imaged every 10 minutes (20 z-slices (20 micron) per image). The full movie was cropped in x and y, and then a maximum intensity projection (MIP) was made. KIF2A-GFTS is labeled in green and SiR-tubulin in magenta. Note that in the single SiR-tubulin channel (left panel) the signal was artificially increased for each individual image to be able to visualise SiR-Tubulin for as long as possible. In the merged panel (middle) the SiR-tubulin (magenta) was not enhanced. Yellow arrows indicate two ICB MTs, one of which resolves before the end of the movie.

*Movie S4*

LSFM-based time lapse imaging of non-treated *Kif2a*$^{GFTS}$ mESCs, incubated with SiR-tubulin. mESCs were imaged every 10 minutes. The full movie was cropped in x and y, and then a maximum intensity projection (MIP) was made. Left panels show the actual movie, right panels show intensity profile (y-axis: intensity value, x-axis: length of profile (pixels)).

*Movie S5*

LSFM-based time lapse imaging of dTAG-13-treated *Kif2a*$^{GFTS}$ mESCs, incubated with SiR-tubulin. mESCs were imaged every 10 minutes. The full movie was cropped in x and y, and then a maximum intensity projection (MIP) was made. Left panels show the actual movie, right panels show intensity profile (y-axis: intensity value, x-axis: length of profile (pixels)).

*Movie S6*
*In vitro* microtubule reconstitution assay performed in the presence of oxygen scavenger, with tubulin labeled in red, the MT seed labelled in magenta, and 3 nM GFP-KIF2A in green. Minus (-) and plus (+) end of the seed are indicated. Yellow rectangle indicates area of accumulation of GFP-KIF2A at the MT minus-end. Movie was acquired for 4 minutes (480 frames, 2 frames per second (fps)) and is played at 20 fps. Note that the MT seed was only imaged for one frame at the beginning of the time lapse experiment. This image is present throughout the movie in the upper panel.

*Movie S7*
LSFM-based time lapse imaging of *Kif2a*$^{GFTS}$ mESCs incubated with SiR-tubulin.The mESC colony was imaged every 10 minutes (25 z-slices (25 micron) per image). The full movie was cropped in x and y, and then a maximum intensity projection (MIP) was made. KIF2A-GFTS is labeled in green and SiR-tubulin in red.
